# Genomic asymmetry of the *Brassica napus* seed: Epigenetic contributions of DNA methylation and small RNAs to subgenome bias

**DOI:** 10.1101/2020.09.08.287995

**Authors:** Dylan J. Ziegler, Deirdre Khan, Nadège Pulgar-Vidal, Isobel A.P. Parkin, Stephen J. Robinson, Mark F. Belmonte

## Abstract

Polyploidy has predominated the genetic history of the angiosperms, and allopolyploidy is known to have contributed to the vast speciation of flowering plants. *Brassica napus*, one of the world’s most important oilseeds, is one such polyploid species originating from the interspecific hybridization of *Brassica rapa* (A^n^) and *Brassica oleracea* (C^n^). Nascent amphidiploids must balance progenitor genomes during reproduction, though the role of epigenetic regulation in subgenome maintenance is unknown. The seed is the pivotal developmental transition into the new sporophytic generation and as such undergoes substantial epigenetic modifications. We investigated subgenome bias between the A^n^ and C^n^ subgenomes as well as across syntenic regions by profiling DNA methylation and siRNAs characteristic of *B. napus* seed development. DNA methylation and siRNA accumulation were prevalent in the C^n^ subgenome and most pronounced early during seed morphogenesis. Hypermethylation during seed maturation was most pronounced on non-coding elements, including promoters, repetitive elements, and siRNAs. Methylation on siRNA clusters was more prevalent in syntenic regions of the C^n^ subgenome and implies selective silencing of genomic loci of the seed. Together, we find compelling evidence for the asymmetrical epigenetic regulation of the A^n^ and C^n^ subgenomes of *Brassica napus* across seed development.

## Introduction

Seed development of *Brassica napus* L. (canola) is a complex yet elegant chapter of the plant lifecycle that starts with the unfertilized ovule and ends with formation of the dry seed (Figure 1A). The first phase of seed development results from a double fertilization event that leads to the establishment of the embryo and the endosperm surrounded and protected by the maternally derived seed coat (Guignard, 1902; Dresselhaus and Franklin-Tong, 2013). Early in the morphogenesis phase, the globular (GLOB) embryo begins to differentiate through programmed cell divisions and establish tissue identities (Tykarska, 1976; Smith and Long, 2010). By the HEART stage, the shoot and root apical meristems organize and will eventually give rise to the shoot and root systems of the next sporophytic generation of the plant (Tykarska, 1979; also reviewed in the model *Arabidopsis* in Jenik et al., 2007). The onset of seed maturation begins with substantial cellular proliferation and growth of the embryo to fill the central cavity of the seed, followed by an arrest of proliferation and the accumulation of storage reserves, leading to the mature green (MG) stage of seed development (Tykarska, 1980; Leviczky et al., 2019). Maturation concludes with the rapid desiccation of the seed in preparation for dormancy at the dry seed (DS) stage.

**Figure 1.**
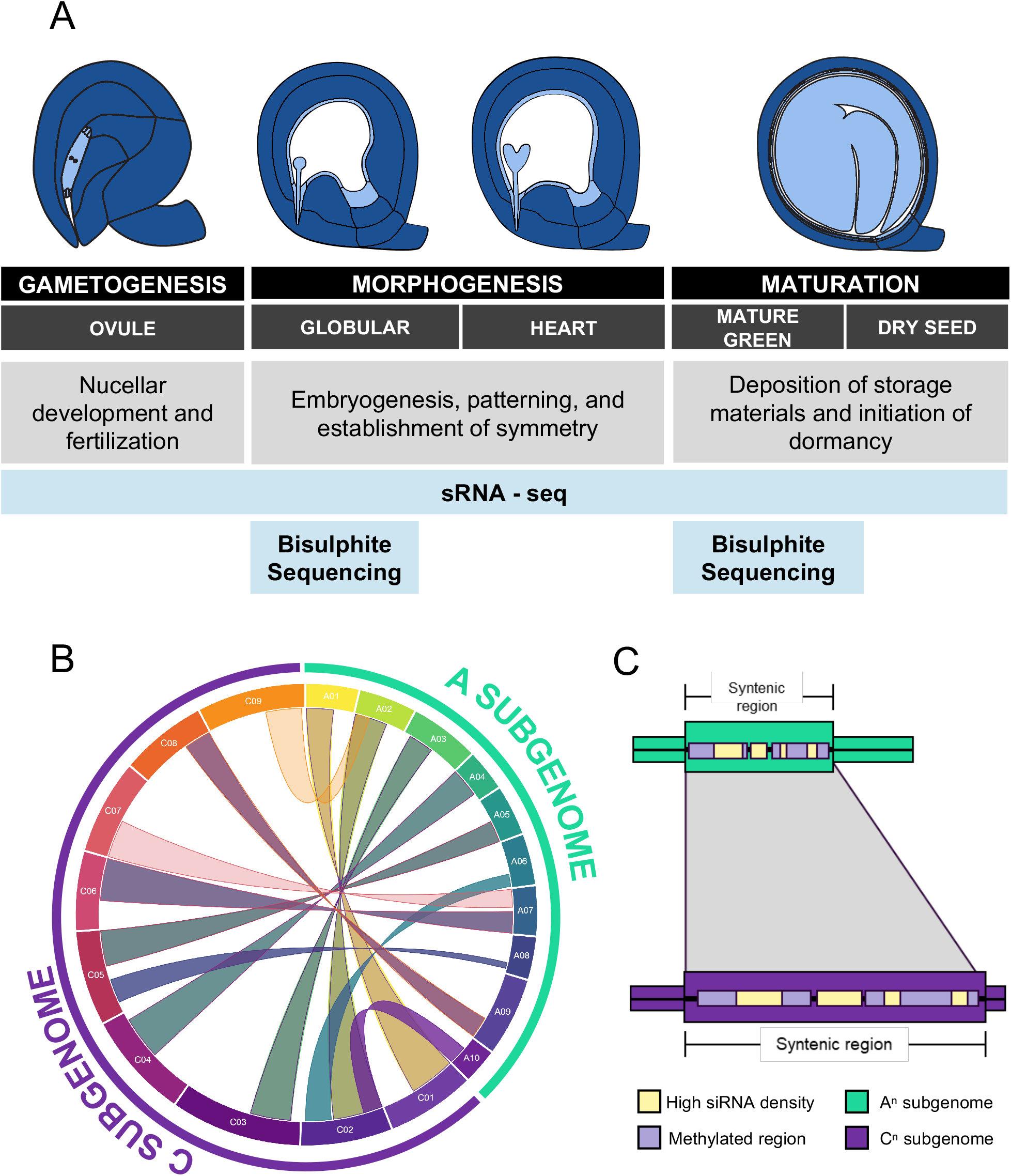
Hypothetical model of epigenetics underpinning seed development in the *B. napus* seed. **(A)** Seed development can be divided into five discrete stages from the beginning of gametogenesis to the end of maturation (ovule (OV), globular (GLOB), heart (HEART), mature green (MG), and dry seed (DS)). We completed sRNA-seq for all five of these stages and bisulphite sequencing for the GLOB and MG stages, representing pivotal developmental transitions in the initiation of morphogenesis and maturation respectively. **(B)** Circos diagram showing the largest continuous syntenic block of each chromosome within the *B. napus* genome. **(C)** Hypothetical model of epigenetic conservation – broad syntenic regions of the genome indicate high conservation between the two subgenomes. Conservation of these regions may lead to similar epigenetic architecture in reproduction.

*B. napus* occupies a unique position in the angiosperm history of polyploidy. While the species shares the ancestral whole genome duplications (WGDs) with *Arabidopsis* that paved the landscape for the diversity of the dicot lineages*, Brassica* is also preceded by a whole genome triplication (WGT) event that defines the Brassiceae (Ren et al., 2018; Franzke et al., 2011). *B. napus* has two discrete diploid genomes from progenitor species *Brassica rapa* (A^r^A^r^) and *Brassica oleracea* (C^o^C^o^), forming the amphidiploid *B. napus* (A^n^A^n^C^n^C^n^). As a result, *B. napus* has a 6x duplication of genetic material relative to *Arabidopsis* and was formed 7500-12500 years ago (Chalhoub et al., 2014). This combination of complicated genomic history with recent speciation means the *B. napus* genome possesses both broad syntenic regions and reshuffling of its progenitor subgenomes (Figure 1B) (Chalhoub et al., 2014; Cai et al., 2014). Thus, *B. napus* represents a valuable plant system for studying genome structure of recent polyploids. Unfortunately, epigenetic phenomena are a poorly understood component of allopolyploid genomes. While transcriptomic data implying subgenome bias in *B. napus* is becoming prevalent in the literature (Wu et al., 2018; Chalhoub et al., 2014), less consideration has been given to other elements of the genome that affect gene expression like DNA methylation or small RNAs (sRNAs). The long-term consequences of subgenome bias are extensive genome fractionation characterized by gene loss and eventual loss of repetitive elements (REs) (Wendel et al., 2018; Cheng et al., 2018, 2014), but the immediate effect allopolyploidization has on the epigenetic landscape, particularly in reproduction, is unknown.

DNA methylation in plants occurs in every cytosine context (CG, CHG, CHH, where H = A, T, or C), with each context exhibiting notably different degrees of methylation and pathways of establishment and maintenance (Cokus et al., 2008; Matzke and Mosher, 2014). Previous works have profiled the DNA methylation signatures during seed development in *Arabidopsis*, maize, *Ricinus communis*, *Oryza sativa*, and *Glycine*, and ovule development in *Gossypium* (Bouyer et al., 2017; Narsai et al., 2017; Xing et al., 2015; Song et al., 2015; Davis-Richardson et al., 2016; Lin et al., 2017). However, studies of DNA methylation in allopolyploid plants such as *Gossypium hirsutum*, *Glycine dolichocarpa*, *Triticum*, and *Erythranthe perigrinus,* has primarily focused on vegetative tissues (Song et al., 2015; Coate et al., 2014; Gardiner et al., 2015; Li et al., 2014, 2017; Chalhoub et al., 2014; Bird et al., 2019; Li et al., 2016; Edger et al., 2017), and thus our understanding of how DNA methylation dynamics compare between progenitor subgenomes in allopolyploids during reproduction and seed development is less well-studied. Furthermore, no current studies have endeavoured to study DNA methylation patterns of seed development in the nascent allotetraploid *B. napus*.

Small RNAs also play a role in the regulation of genetic material encoded by the plant genome. The most studied sRNAs, micro RNA (miRNA), are involved in the regulation of gene expression via post-transcriptional silencing. miRNAs arise from a single stranded precursor which forms a secondary hairpin structure after transcription (Wang et al., 2019). Unlike miRNAs, small interfering RNAs (siRNA) do not solely operate in post-transcriptional silencing and can induce changes within the genome itself. siRNAs differ from miRNA in their mode of biogenesis, wherein siRNAs are transcribed by RNA Pol IV/V, and a complementary strand is synthesized by RNA-dependent RNA polymerase (RDR) to form a double-stranded siRNA precursor (Zhou and Law, 2015). siRNAs can be encoded by both intergenic and intronic regions, but can also originate from REs including transposable elements (TEs) and heterochromatic regions (Chen et al., 2018). siRNAs are capable of transcriptional silencing like miRNAs via natural anti-sense siRNA (NAT-siRNA) but can exert more versatile functions in genomic regulation. Repeat associated siRNA (rasiRNA), referred to in some literature as heterochromatic siRNA, are typically encoded by REs and are most frequently associated with the silencing of TEs via RNA-directed DNA methylation (Wang and Axtell, 2017). Phased siRNA (phasiRNA) are encoded by a phased locus with repeating segments that are cleaved into multiple 21- or 24-nt mature phasiRNAs principally by DCL4 in dicots (Deng et al., 2018). siRNAs contributing to plant development are surfacing in the literature, but few consider siRNAs in the context of allopolyploid genomic bias. Our knowledge on how siRNAs accumulate in subgenomes of amphidiploids and the presence of subgenome bias is currently limited to vegetative tissues, with minimal information on these dynamics in reproduction (Shen et al., 2015). It is also unknown if certain classes of siRNAs accumulate asymmetrically or if these siRNAs are ancestrally conserved within subgenomes. To date, research on siRNA populations of *B. napus* are scarce and fewer profile siRNAs occurring across seed development.

The orchestration of seed development relies on a number of genomic factors. It is known that DNA methylation and RNA interference are critical processes underpinning development, and most studies examining patterns of methylation and non-coding RNA accumulation in the seed are in the model Arabidopsis (D’Ario et al., 2017; Bouyer et al., 2017; Kirkbride et al., 2019; Narsai et al., 2017). To date, little is known about epigenetics and their dynamics between subgenomes during seed development in amphidiploids like *B. napus*. To address these questions, we profiled both the DNA methylation and siRNA landscape across seed development and at critical developmental transitions of the seed lifecycle. Mapping of syntenic regions across the two progenitor subgenomes established horizontal comparisons between the A^n^ and C^n^ subgenomes (Figure 1C). Taken together, we find strong evidence for C^n^ subgenome bias in the context of both DNA methylation and siRNA diversity, and find that the A^n^ subgenome exerts epigenetic dominance over the submissive C^n^ subgenome in seed development.

## Results

### A and C subgenomes share vast syntenic regions

Broad syntenic blocks are observed in both A^n^ and C^n^ subgenomes of *B. napus* (Figure 1B). Here, we define syntenic regions as genomic blocks of colinear genes shared between the A and C subgenomes and recognize that the chromosomes of the A^n^ and C^n^ subgenomes are not entirely homologous in structure *sensu stricto.* As a result, we consider any blocks of colinear genes to be syntenic regions regardless of where the matching syntenic region is within the other subgenome. We report 778 syntenic regions spanning the A^n^ and C^n^ subgenomes collectively (>10kb) and account for 99.624% of the genome assembly (Dataset S1). Chromosomes are not horizontally comparable between the A^n^ and C^n^ subgenomes. While some chromosome pairs exhibit robust homology (ie. A01 and C01), others are more fractionated, with homologous regions largely spread throughout the other subgenome (ie. A06 and C06; Dataset S1). We used this information to frame the analysis of the A^n^ and C^n^ subgenomes in the context of DNA methylation and siRNA diversity defining *B. napus* seed development.

### The epigenetic landscape of seed development

Despite the high degree of synteny between the A^n^ and C^n^ subgenomes, there is a notable difference in structure between the two subgenomes of *B. napus* cv. DH12075. The smaller A^n^ subgenome had a higher density of gene models than the C^n^ subgenome (Figure 2A). The larger Cn subgenome had accumulated a greater density of repetitive elements (REs) that were also larger in size than in the A^n^ subgenome (A^n^: 192,038 REs, C^n^: 293,260 REs; Figure 2A; Figure S1; F-Test, p<0.001; A^n^ RE average = 396bp, C^n^ RE average=608bp). In both subgenomes, mCG and mCHG was focused on areas of high RE density (Figure 2B). This corresponded with larger swathes of highly methylated cytosines in the RE-rich C^n^ subgenome (Figure 2B, Table 1, Dataset S2). By contrast, active siRNA loci were distributed in areas of low RE density and high gene density (Figure 2C). An increase in CHH methylation was observed at the MG stage of seed development (Table 1, Dataset S2). Correspondingly, there was a genome-wide burst in siRNA detection at the MG stage of development (Figure 2C).

**Figure 2.**
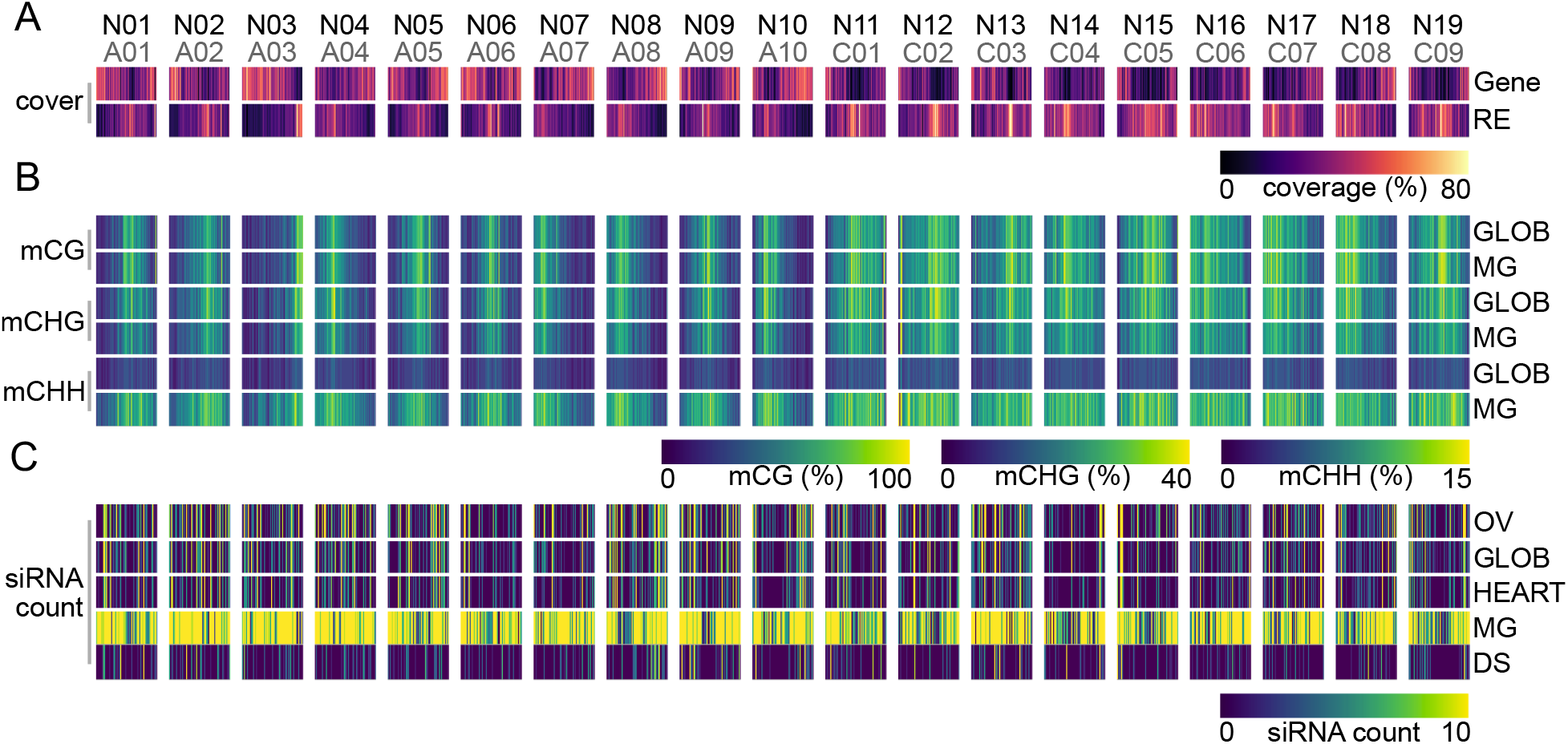
The epigenetic landscape of the *Brassica napus* seed. Both the N (black) and A/C (grey) chromosome notation are indicated for chromosomes (A) Coverage (%) of 100kB fixed windows by genes and repetitive elements (RE) across the DH12075 *B. napus* genome. High coverage (80%) is indicated by yellow fill, and low coverage (0%) is indicated by dark purple fill. (B) Average methylation in each cytosine context (CG, CHG, CHH) are shown separately for the GLOB and MG stages of seed development. High methylation levels (CG: 100%, CHG: 40%, CHH: 15%) are indicated by a yellow colour, and low methylation (0%) is indicated by a purple fill. (C) Total number of detected/active siRNA clusters in 100kB bins across the ovule (OV), globular (GLOB), heart (HEART), mature green (MG), and dry seed (DS) stages of development. A high count (10) is indicated by yellow fill, and low counts (0) are indicated by purple fill.

### Single base pair resolution profiling of the *B. napus* seed methylome reveals C^n^ subgenome bias

Genome-wide cytosine methylation was calculated over 1kB windows for both GLOB and MG seed stages in all cytosine contexts and was compared to available methylation data for leaves (Table 1, Dataset S2). We estimated bisulfite conversion efficiency using the unmethylated plastid genome as a reference (Ji et al., 2014), and estimated the bisulfite conversion rate to be ~98% in both the GLOB and MG samples (Figure 3A). We compared GLOB and MG seed methylome data to leaf methylome data, as a developmental comparison to a differentiated, vegetative tissue. Relative to leaves, seed development was characterized by a significant loss of mCG and mCHG levels (Table 1, p<0.001 Mann-Whitney-Wilcoxon). However, mCHH levels were nearly 3-fold higher at the MG stage than observed in globular seeds or in leaves (Figure 3A, Table 1, p<0.001 Mann-Whitney-Wilcoxon). This increase in mCHH contributed to a 2.5% increase in overall cytosine methylation of MG compared to GLOB seeds (Figure 3A).

**Figure 3.**
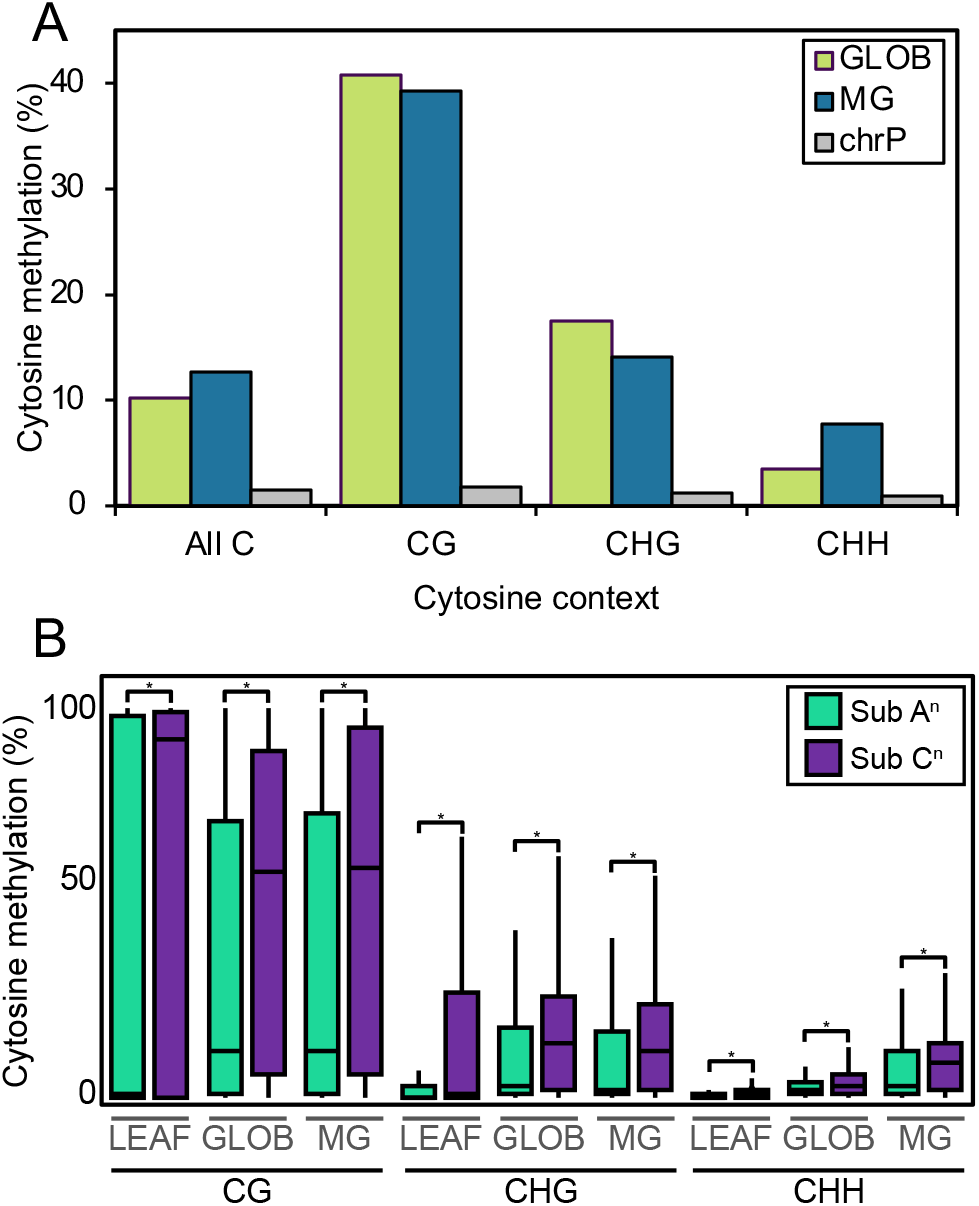
The DNA methylation in *Brassica napus* seed development. (A) Average methylation levels by cytosine context. Average methylation for all covered cytosines is shown for the GLOB (green), MG (blue), and plastidial (grey) genomes. Level of methylation in the largest syntenic regions between the A^n^ (teal) and C^n^ (purple) subgenomes of *B. napus*. Quantile boxplots show level of cytosine methylation as calculated in 1kB windows in the syntenic regions for leaves, and GLOB and MG seeds in the CG, CHG, and CHH contexts. Significant differences in cytosine methylation between the two subgenomes is indicated with an asterisk (p<0.001 Mann-Whitney-Wilcoxon, Bonferroni adjustment for multiple comparisons).

We further identified differences in DNA methylation between leaves, GLOB, and MG seeds in addition to significant differences in the methylation of the A^n^ and C^n^ subgenomes of *B. napus*. In leaves, GLOB and MG seeds, mCG and mCHG levels were 1.4-1.5x higher in the C^n^ than in the A^n^ subgenome (Table 1, Mann-Whitney-Wilcoxon, p<0.001). Significant subgenome bias was also observed in CHH methylation, though mCHH levels were 1.1-1.2x higher in the C^n^ subgenome than in the A subgenome. A sharp increase in mCHH during seed maturation nearly abolishes mCHH subgenome bias in MG seeds (Table 1, Mann-Whitney-Wilcoxon, p<0.001). Overall, while subgenome methylation bias persisted in *B. napus* leaves and seeds, this phenomenon was strongest in leaves and weakens in the seed as it matures (Table 1). We also compared cytosine methylation between the largest syntenic blocks between the A^n^ and C^n^ subgenomes (Figure 3B, Dataset S3), and found that a bias towards higher methylation of the C^n^ subgenome was maintained over these high homology regions. However, there was variation in the extent of subgenome bias between the A^n^ and C^n^ syntenic blocks. CG and CHG methylation were only 1.2x higher on the C^n^ counterpart in the A04/C04 syntenic pair, but was as high as 3.2x (CG) - 3.3x (CHG) higher on the C^n^ counterpart on the A08/C05 pair in GLOB and MG seeds. There is a positive trend between degree of C^n^/A^n^ subgenome bias with an increased relative C^n^/A^n^ syntenic block size (R^2^ = 0.70-0.78 depending on cytosine context; Dataset S1). This may be a result of increased RE density in the C^n^ subgenome (Figure 2A) as REs exhibit high methylation in all contexts (Figure 5A, Table 2).

### Unique suites of promoters are differentially methylated targeting genes involved in development and carbon metabolism

Methylation on promoters is relatively stable during seed development. CG methylation on promoters does not significantly change between GLOB and MG seed stages (leaf mCG = 71.8, GLOB mCG = 64.0, MG mCG = 63.8; Figure 4A; Table 2; p<0.001 Mann-Whitney-Wilcoxon). When GLOB and MG seeds were compared to leaves, 84.6% of CG differentially methylated promoters, and 77.8% of CHG differential methylation on promoters, was shared between the GLOB and MG seeds (Figure 4B, Figure S2), suggesting stability in promoter CG methylation patterns once seed development has initiated. Combined, these data suggest the developing seed is distinguished from vegetative leaves by lower levels of CG and CHG methylation on promoters that is maintained through the development and maturation of the seed. Subgenome bias in DNA methylation was also maintained through this shift. While CG methylation was significantly higher on GLOB and MG C^n^ subgenome promoter elements (Figure 4A; Table 2), we did not identify significant differences between the A04/C04 and A07/C07 high homology regions (Dataset S3). Additionally, there was no significant difference in CHG methylation on the A04/C04, A05/C05, and A07/C07 syntenic regions despite an overall bias towards higher CHG methylation of the C^n^ subgenome promoters (Figure 4A; Table 2, Dataset S3).

**Figure 4.**
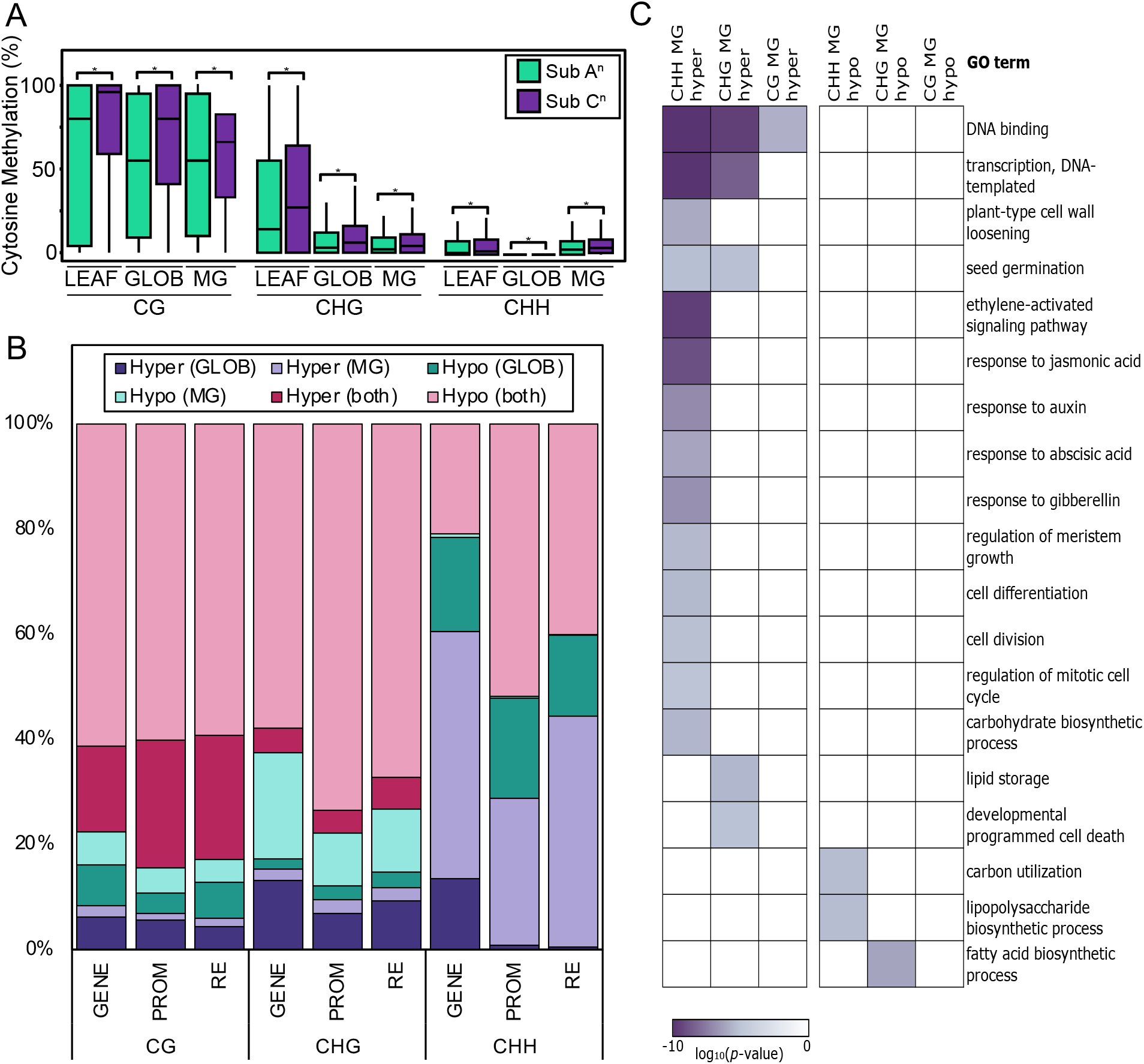
Shifts in DNA methylation between in development. (A) Quantile boxplots show level of cytosine methylation on promoters (1kb upstream of TSS) in the syntenic regions for leaves, and GLOB and MG seeds in the CG, CHG, and CHH contexts. Significant differences in cytosine methylation between the A^n^ (teal) and C^n^ (purple) subgenomes is indicated with an asterisk (p<0.001 Mann-Whitney-Wilcoxon, Bonferonni adjustment for multiple comparisons). (B) Stacked 100% column showing hyper- and hypo-methylated genes (GENE), 1kB upstream regulatory regions (promoters, PROM), and repetitive elements (RE) between seeds (GLOB and MG) and leaves. Data are separated into hyper-methylated in GLOB (dark purple) or MG seeds (light purple), hypo-methylated in GLOB (dark teal) or MG (light teal) seeds, and hyper-methylated (dark pink) or hypo-methylated (light pink) in both GLOB and MG stages. (C) GO enrichment of genes with differentially methylated promoters between the GLOB and MG stages of seed development. Highly enriched GO terms are indicated by darker colouring. Gene lists and enrichment output can be found in Dataset S4.

There is a significant increase in CHH methylation on promoter elements during seed development which favours higher mCHH on C^n^ subgenome promoters (Figure 4A) in the A03/C03, A07/C06, and A02/C09 syntenic regions (Table 2; GLOB mCHH = 0.5%, MG mCHH = 6.4; p<0.001 Mann-Whitney-Wilcoxon; Dataset S3). We compared promoter element methylation of seeds to leaves and found that of the 1976 promoters differentially methylated between GLOB (933) and MG (1043) seeds and leaves (Figure 4B; Figure S2), 95% were derived from the C^n^ subgenome (Dataset S4). To identify the biological processes affected by developmental shifts in promoter methylation, we examined enrichment of Gene Ontology (GO) terms for genes downstream of promoters differentially methylated between GLOB and MG seeds (Figure 4C, Dataset S4; significance of GO enrichment considered at p<0.001). At seed maturation, predicted biological processes involved in gene activation were heavily enriched in genes downstream of hypermethylated promoters, such as DNA binding (CG, CHG, CHH hypermethylated) and DNA-templated transcription (CHG and CHH hypermethylated). CHH hypermethylation of promoters in MG seeds was also enriched for response to hormones such as ethylene, jasmonic acid, gibberellin, and auxin. We also observed CHH hypermethylation of promoters upstream of genes involved in cell division and differentiation, cell wall loosening, programmed cell death, and seed germination in GLOB seeds. Promoter hypomethylation of MG seeds were enriched for fatty acid biosynthesis (CHG hypermethylation) and carbon utilization (CHH hypermethylation), while promoters of genes involved in carbohydrate biosynthesis were targeted for hypermethylation (CHH). Therefore maturation-phase promoter hypermethylation appears to target hormone response, gene activation, and the balance between morphogenesis and quiescence, while promoter hypomethylation in seed maturation is likely associated with the release of genes for mobilizing carbon into lipid biosynthesis (Figure 4C).

### Transcription factors are hypomethylated relative to other protein-coding genes

We examined the methylation levels of transcription factor (TF) genes relative to other protein coding genes and found significantly lower methylation on gene bodies and promoters of TF-coding genes (Figure S3; Table 2; p<0.001 Mann-Whitney-Wilcoxon). In leaves, GLOB, and MG seeds, transcription factor gene body methylation levels were reduced by 57-78% relative to the levels of other coding gene bodies (Figure S3; Table 2, p<0.001 Mann-Whitney-Wilcoxon). Transcription factor promoter CG and CHG methylation is significantly reduced relative to other coding genes, though reduction is less than observed on gene bodies. While mCG on seed TFs was significantly lower than observed in leaves, mCG levels did not significantly change between GLOB and MG seeds nor did they change on the promoters of TFs (Table 2; p<0.001 Mann-Whitney-Wilcoxon). Additionally, subgenome methylation bias was maintained on TF gene bodies and promoter regions (Table 2).

### Methylation bias on repetitive elements during seed development

In all three cytosine contexts, the vast majority of differential methylation between leaf and seed methylomes occurred on repetitive elements (Figure S4, Dataset S4). Methylation of CG and CHG contexts on REs was significantly reduced in seed development relative to leaves (leaf mCG = 91.5%; Table 2). CG methylation on REs was observed to be lower during seed morphogenesis and increased during seed maturation (GLOB mCG = 80.4%, MG mCG = 84.6%; Table 2; significant p<0.001 Mann-Whitney-Wilcoxon). Overall, a degree of subgenome bias was maintained during these shifts in CG methylation: mCG levels are 1.1x higher on C^n^ subgenome REs (Table 2; p<0.001 Mann-Whitney-Wilcoxon); higher methylation of the C^n^ subgenome REs is also observed between the largest syntenic blocks between the A^n^ and C^n^ subgenomes (Figure 5A). Further comparison of the REs within the largest syntenic regions between A^n^ and C^n^ subgenomes shows significant difference at the MG stage of seed development on A03/C03, A07/C06, and A02/C09 high homology regions (Dataset S3). Therefore, bias towards higher methylation of C^n^ subgenome REs is most prevalent during seed maturation and is not evenly distributed throughout the genome.

**Figure 5.**
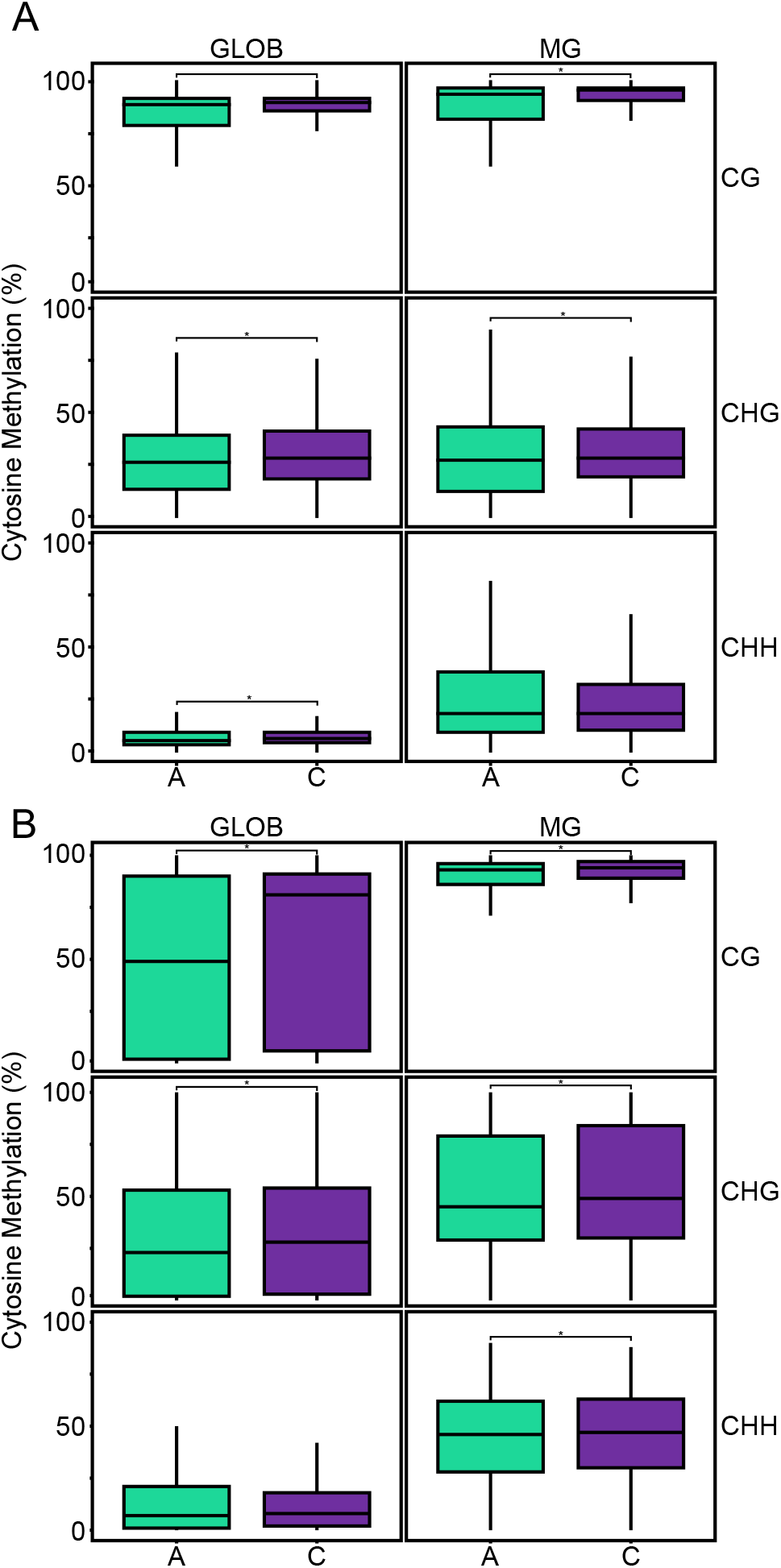
Subgenome methylation bias on repetitive elements and siRNAs. (A) Quantile boxplots showing the methylation levels of repetitive elements (A) and active/detected siRNA clusters (B) in the largest syntenic regions between the A^n^ (teal) and C^n^ (purple) subgenomes for GLOB and MG seeds in the CG, CHG, and CHH cytosine contexts. Significant difference of methylation levels between the two subgenomes is indicated by an asterisk (*) (Mann Whitney Wilcoxon, p<0.001, Bonferonni correction for multiple testing).

CHG methylation on REs does not change significantly between the GLOB and MG stages of seed development (GLOB mCHG = 30.8%, MG mCHG = 32.4%, Table 2; p<0.001 Mann-Whitney-Wilcoxon; Figure 5A). However, there is significantly higher CHH methylation on a unique suite of REs in maturation that is observed neither in leaves nor during seed morphogenesis (Figure S2). Lower of CG and CHG methylation relative to leaves is countered by an increase in CHH methylation on REs during seed maturation (leaf mCHH = 9.6%, GLOB mCHH = 7.2%, MG mCHH = 23.0%; Table 2; all significant p<0.001 Mann-Whitney-Wilcoxon) and is observed globally as well as within the largest syntenic regions of the genome (Figure 5A). Over 40% of CHH-DM REs were hypermethylated in the MG stage relative to leaves (Figure S2). CHH methylation of A^n^ subgenome REs was 1.2x higher than C^n^ subgenome REs in leaves, though this bias was abolished in the seed (Table 2, p<0.001, Mann-Whitney-Wilcoxon), especially during the MG stage wherein comparison of the largest syntenic regions reveals no mCHH subgenome bias on REs (Figure 5A). This suggests extensive reprogramming of genome architecture around REs during seed development, driven by both a loss of CG and CHG methylation, and subgenome-independent increase of CHH methylation on REs during seed maturation.

### Methylation of SRCs during seed maturation

Cytosine methylation increased 1.5-3.5 fold on siRNA clusters (SRCs) from the GLOB to the MG stage of seed development (GLOB mCG = 54.4%, MG mCG = 84.6%; GLOB mCHG = 33.0%, MG mCHG = 52.22%; GLOB mCHH = 12.5%7, MG mCHH = 44.08%; Table 2; p<0.001 Mann-Whitney-Wilcoxon). Most notably, the mCHH levels of SRCs at the MG stage are 4.9x higher than the whole genome average (Table 2), and nearly double the levels observed on REs (Figure 5A-B).

In general, we did not observe significant methylation bias on siRNAs between the A^n^ and C^n^ subgenomes (Table 2). However, in the largest syntenic blocks, a bias towards higher C^n^ subgenome methylation in all cytosine contexts in GLOB and MG seeds was observed, with the exception of mCHH in GLOB seeds (Figure 5B). There was also a bias towards higher CG methylation of C^n^ subgenomes in the A02/C02 (MG), A02/C09 (MG), A03/C03 (GLOB and MG), A04/C05 (MG), A07/C06 (GLOB and MG), A08/C05 (GLOB and MG) syntenic blocks (Dataset S3). In the CHH context, where the most dramatic shift in SRC methylation was observed, we identified bias on clusters in the A06/C02 and A08/C05 syntenic blocks (Dataset S3). Therefore, bias in the epigenome with regards to siRNAs is limited to comparisons within syntenic regions.

### siRNAs undergo shifts in diversity across seed development

Next, we profiled small RNA populations, genome wide across *B. napus* seed development. The most striking differences occurred in the diversity of active SRCs across the five stages of seed development outlined in this work. For example, the MG stage of seed development contained the most active SRCs (61,929, Figure 6A). Of them, 41% and 59% were encoded by the A^n^ and C^n^ subgenome, respectively. This trend of bias towards the C^n^ subgenome persists across all other seed stages, with 23,312 SRCs in the OV stage (39.7% and 60.3%), 18,541 in the GLOB stage (41% and 59%), 10,276 in the HEART stage (42% and 58%), and 4,800 in the DS stage (41.5% and 58.5%). The DS exhibits less than half of the total SRCs of the HEART stage, which was otherwise the least diverse stage. A complete list of all SRCs across seed development is found in Dataset S6.

**Figure 6.**
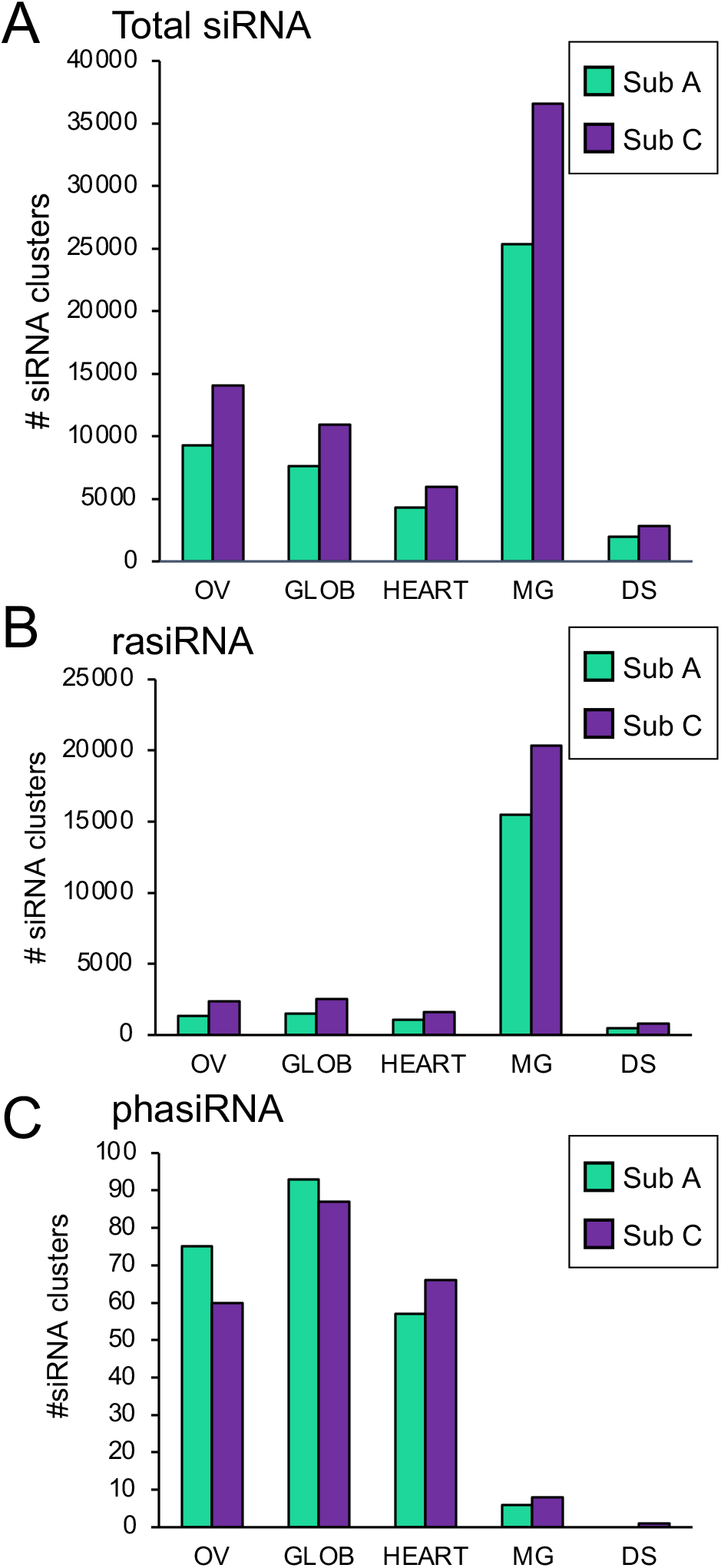
siRNA cluster diversity of five developmental seed stages in *Brassica napus*, beginning in gametogenesis (OV), and ending in maturation and dormancy (DS). **(A)** Total diversity of active siRNA clusters in five seed developmental stages by subgenome (A subgenome in bright green, C subgenome in purple). Number of unique clusters of **(B)** rasiRNA and **(C)** phasiRNA per seed stage.

Active rasiRNA and phasiRNA clusters underwent transitional changes in diversity throughout seed development. rasiRNAs increase in diversity particularly during the MG stage where a total of 35,840 active RNAs were identified (Figure 6B, Dataset S7). The GLOB stage seed exhibited the next-most diverse rasiRNA profile of 4,015 clusters, an order of magnitude less than that of the MG seed. C^n^ subgenome bias was still prevalent in rasiRNA, with total rasiRNA clusters being 64%, 62.6%, 60.3%, 56.8%, and 63% encoded by the C^n^ subgenome in the OV, GLOB, HEART, MG, and DS stages, respectively. rasiRNAs accumulated across seed development, increased in diversity early in maturation, and sharply declined once the seed entered dormancy. rasiRNAs were an abundant siRNA type in our analysis - constituting 15.8%, 21.7%, 26.0%, 57.8%, and 26.4% of the total SRCs in the OV, GLOB, HEART, MG, and DS stages, respectively.

phasiRNA clusters were most diverse early in seed development, with 143, 184, 129, 15, and 1 active cluster(s) in OV, GLOB, HEART, MG, and DS stages, respectively (Figure 6C, Dataset S7). phasiRNAs, however, instead show bias encoding loci towards the A^n^ subgenome in early stages of development, wherein 55.6%, 51.7%, 46.3%, 42.9%, and 0% of the phasiRNA clusters are located within the A^n^ subgenome in OV, GLOB, HEART, MG, and DS, respectively with the DS only having one detected phasiRNA. Interestingly, all mature phasiRNAs were predicted to be 24-nt long aside from the sole cluster in the DS stage, which was 21-nt. Thus, phasiRNAs are the only instance in which SRCs exhibit bias in the A^n^ subgenome, and only in the early stages of reproductive development.

### Promoter regions are implicated in siRNA regulatory mechanisms during seed maturation

We furthered our study of promoter regions and gene bodies by identifying SRCs within them. We find that SRCs spike in diversity during the MG stage in both gene body and promoter contexts (Figure 7A, B). C^n^ subgenome bias was universal across all seed stages and in both gene bodies and promoters. Subgenome bias also exhibited minimal fluctuations throughout development, with the C^n^ subgenome encoding 57.1%, 54.7%, 54.3%, 54.8%, and 54.9% of the SRCs in gene bodies and 58.8%, 57.0%, 55.8%, 55.8%, and 58.0% within promoters in the OV, GLOB, HEART, MG, and DS stages, respectively. The MG stage is characterized by high diversity of SRCs in promoters, especially relative to that in gene bodies. 3295 and 4000 unique SRCs originated from gene bodies in the MG stage, while promoter regions accounted for 6232 and 7872 SRCs in the A^n^ and C^n^ subgenomes, respectively. The disparity between siRNAs encoded in gene bodies and promoters during the MG stage was not noted in any other developmental stage to that level, with all other stages having less SRCs originating from promoter regions than gene bodies. Thus, the dramatic influx of siRNAs from promoter regions during the MG stage provides epigenetic evidence into this pivotal developmental transition of the seed lifecycle.

**Figure 7.**
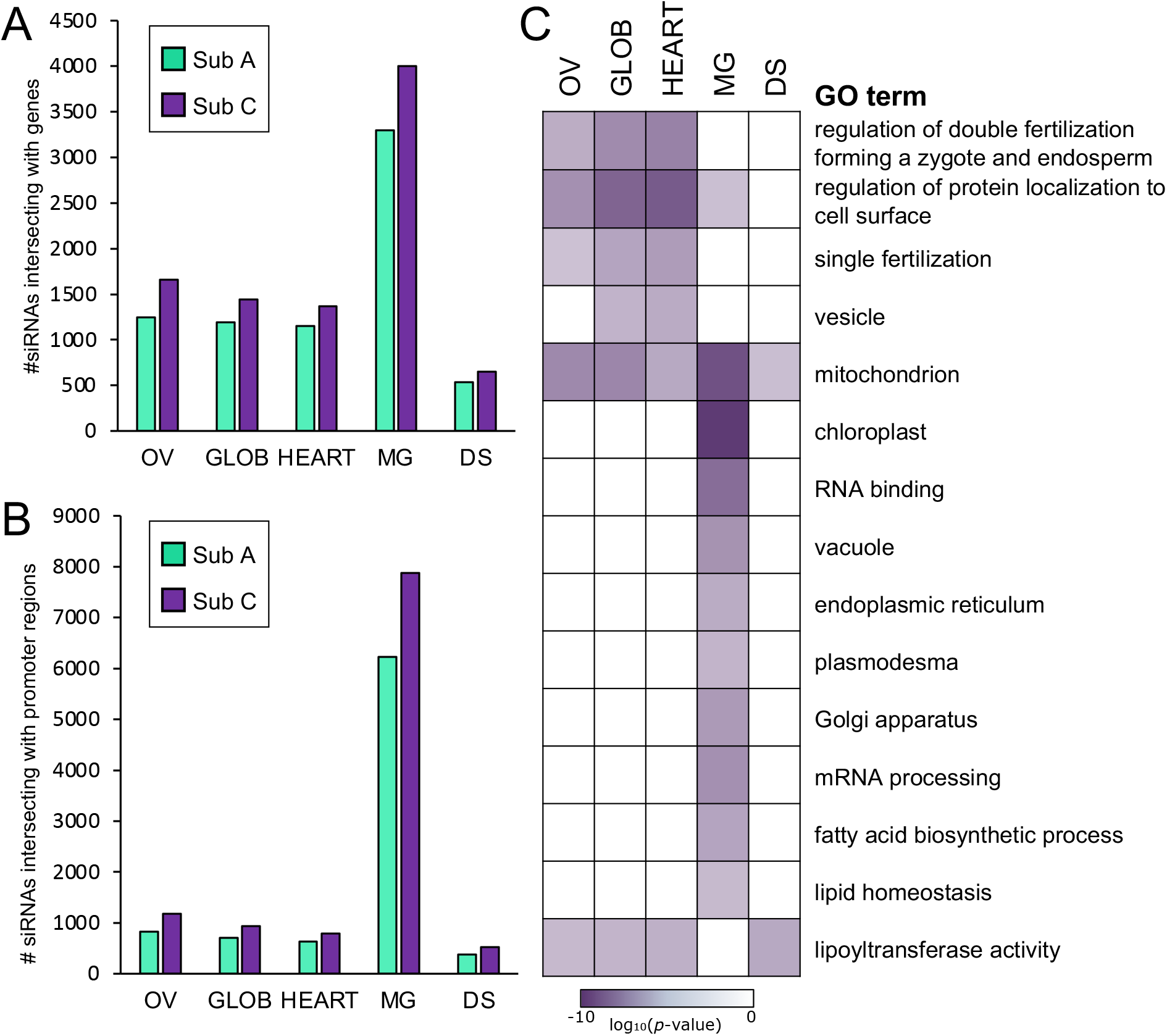
siRNA clusters encoded by gene bodies and promoters. **(A)** Clusters of siRNAs originating from gene bodies spike in the MG stage, with consistent C^n^ subgenome bias. **(B)** Clusters of unique siRNAs encoded within the promoter region of gene bodies also spike in the MG stage, with a notable increase in diversity than seen in gene bodies. **(C)** GO analysis of coding genes associated with the promoter region encoding siRNA clusters

### siRNAs are implicated in developmental processes of the seed

Due to the high siRNA diversity in the promoter context at the initiation of seed maturation, we performed GO enrichment analysis on the genes downstream of the associated SRCs encoded by promoter regions (Figure 7C, Dataset S8). We find that GO terms relating to zygote and endosperm development, protein localization, fertilization, and vesicle formation/transport to be significantly enriched early in seed development (gametogenesis and morhogenesis) (p < 0.001). Though genes associated with mitochondrial activity were significantly enriched at every stage, we find genes associated with chloroplast activity were particularly (p < 0.00001) enriched in the MG stage. Further, GO terms associated with the endomembrane system (endoplasmic reticulum, Golgi apparatus, plasmodesma) were significantly enriched at the MG stage. While both genes associated with fatty acid biosynthesis and lipid homeostasis were enriched in the MG stage, lipoyltransferase activity was instead enriched significantly in every stage except MG. The intervention of siRNAs in the regulation of genes related to cellular organization (endosperm development and protein localization) early in seed development in addition to the crucial biological processes implicated in seed maturation (lipid homeostasis, endomembrane activity) suggest that siRNAs may be playing a yet to be explored role in the maintenance of morphogenesis and maturation.

### Epigenome architecture of the *B. napus* seed is preserved in highly homologous chromosomes

The A01/C01 chromosome pair was over 99% colinear, implying high homology. The two major syntenic blocks on A01 and C01 were 15.7 Mb/26.5 Mb and 9.6 Mb/17.3 Mb in the A^n^/C^n^ subgenomes, respectively (blocks 1 and 2, Figure 8). The DMRs were distributed similarly across all cytosine contexts, with CG and CHG methylation changing comparatively little between the GLOB and MG stages and CHH experiencing dramatic methylation shifts during seed maturation. siRNA density was highest in intergenic regions in both the GLOB and MG stages (Figure 8, Dataset S9). The two largest syntenic blocks (blocks 1 and 2, Figure 8) had similar siRNA densities. For example, siRNAs had the highest density on A01 further from the highly repetitive central region (presumed to be the centromere) and peaked at 65 and 213 unique active SRCs per 100 kb in the GLOB and MG stages, respectively. This trend was also detected in the syntenic region of C01, wherein siRNA density peaked at 55/54 and 208/161 on either side of the centromere in the GLOB and MG stages, respectively. The exception to this trend appears in a steep incline in SRCs in both the GLOB and MG stages on C01, wherein siRNA cluster density increases to 95 and 116 in the region neighboring the centromere. More highly repetitive regions also exhibit fewer changes in CG and CHG methylation contexts from morphogenesis to maturation. Together, this high homology chromosome pair shares similar genomic architecture across seed development and provides evidence into the conservation of epigenetic reprogramming amidst regions in this amphidiploid plant.

**Figure 8.**
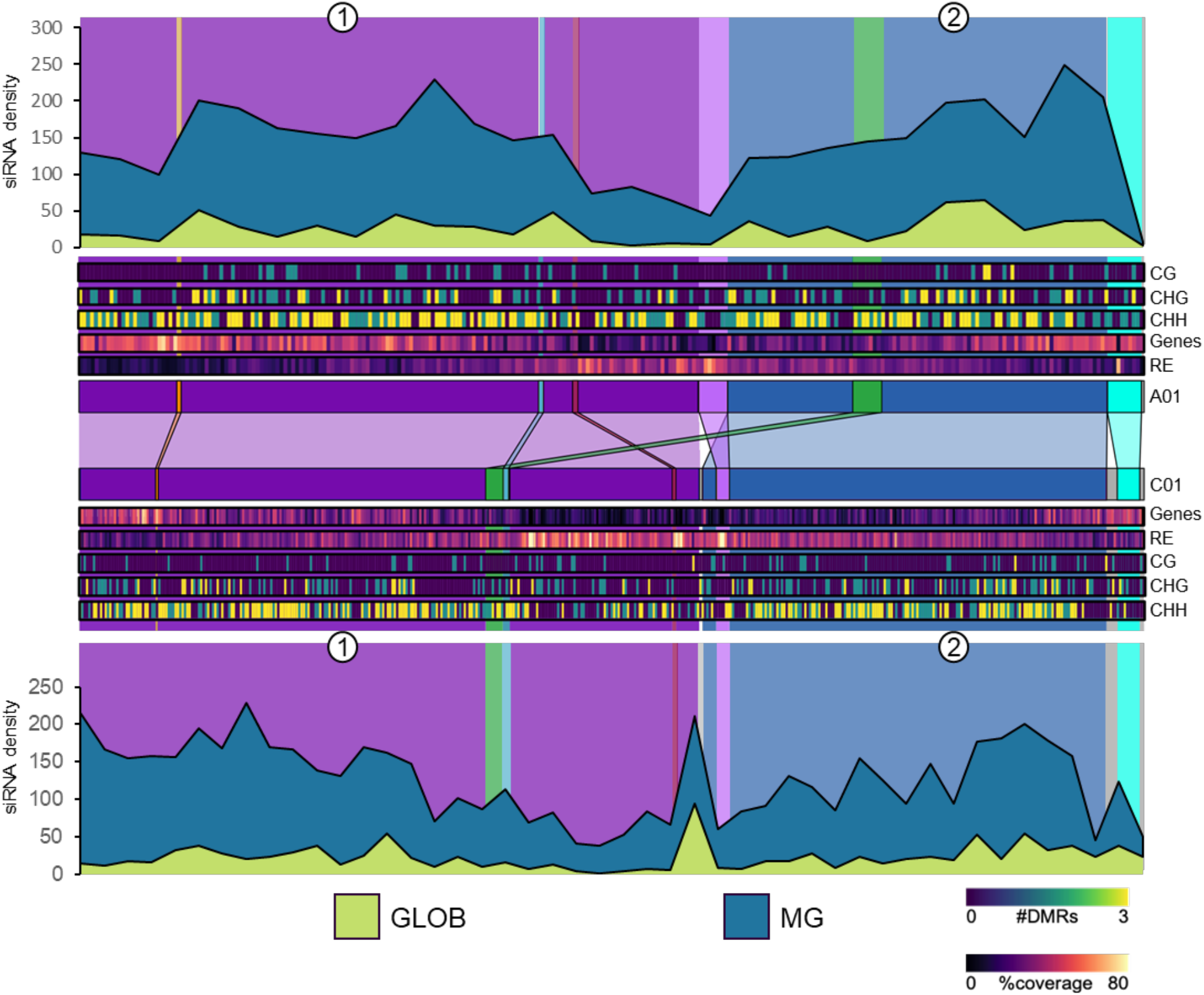
Genomic architecture of a high homology chromosome pair (A01 and C01). Synteny map linking homologous regions between A01 and C01 are located to the centre. Colours on the synteny map represent matching syntenic regions between the A^n^ and C^n^ subgenomes, grey areas indicate no corresponding synteny. Area plots represent siRNA cluster density in the GLOB and MG seed stages in 100kbp windows. Heat maps inadicate both the density of protein-coding genes and repetitive elements (RE), as well as GLOB-MG differentially methylated regions (DMRs) of each cytosine context (CG, CHG, and CHH). Large syntenic regions of interest are marked (1, 2) and referenced in text.

### Low homology chromosomes conserve less epigenetic structure across seed development

We used the A06/C06 chromosome pair as a model for low homology since these two chromosomes have smaller syntenic regions with larger non-syntenic gaps (Figure 9). The largest syntenic blocks were 2.6 Mb/6.2 Mb and 1.3 Mb/1.1 Mb in the A^n^/C^n^ subgenomes, respectively, with many smaller colinear blocks found at 116 kb/190 kb and 219 kb/538 kb (Dataset S1). We compared the syntenic blocks of low homology chromosomes to gain insight into the epigenetic restructuring of polyploid plants across seed development.

**Figure 9.**
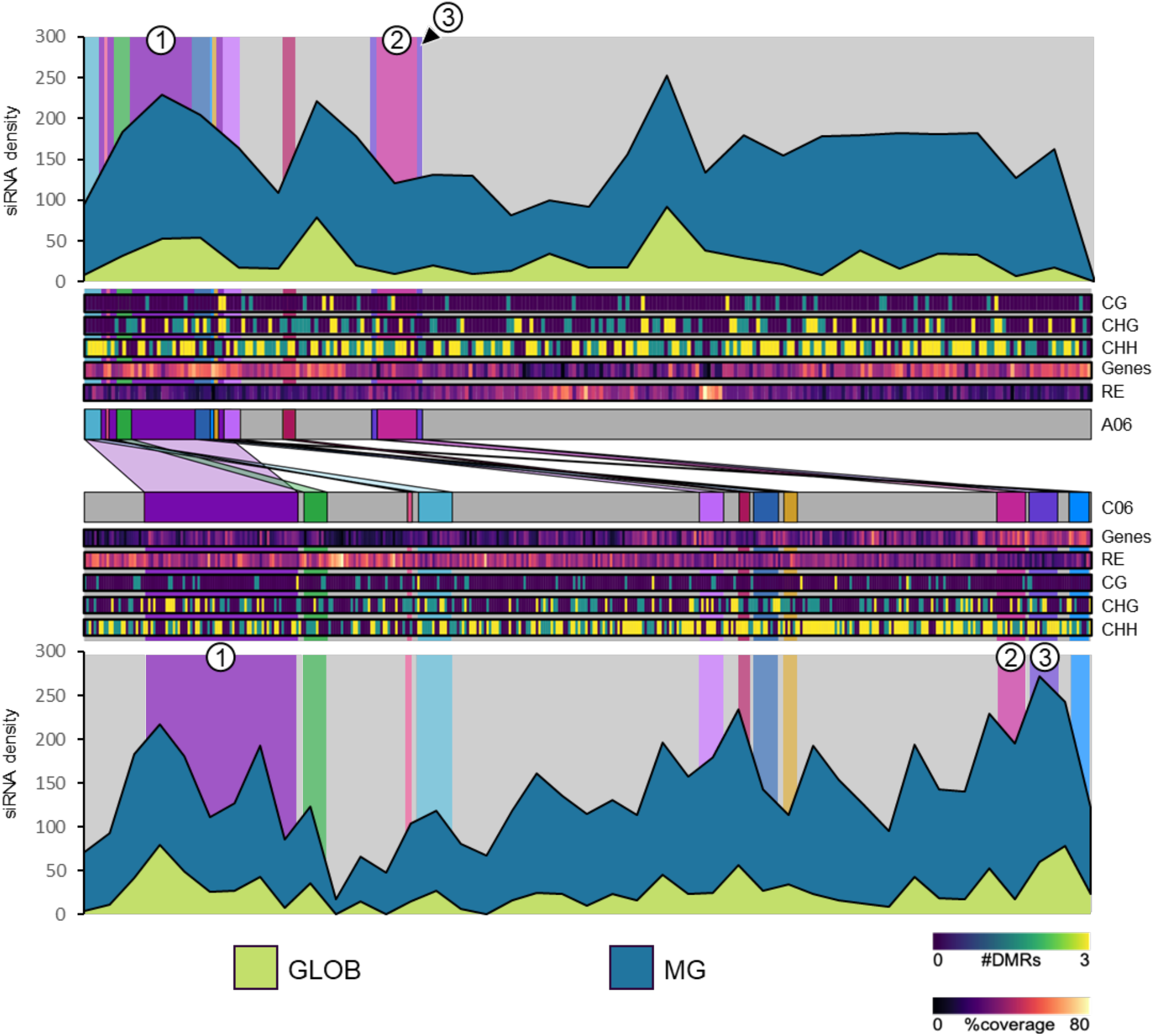
Genomic architecture of a low homology chromosome pair (A06 and C06). Synteny map linking homologous regions between A06 and C06 are located to the centre. Colours on the synteny map represent matching syntenic regions between the A^n^ and C^n^ subgenomes, grey areas indicate no corresponding synteny. Area plots represent siRNA cluster density in the GLOB and MG seed stages in 100kbp windows. Heat maps indicate both the density of protein-coding genes and repetitive elements (RE), as well as GLOB-MG differentially methylated regions (DMRs) of each cytosine context (CG, CHG, and CHH). Syntenic regions of interest are marked (1, 2, 3) and referenced in text.

In the largest syntenic block (block 1, Figure 9), two major peaks of active SRCs were identified in the C^n^ subgenome (79 and 43 active clusters in the GLOB stage, 141 and 150 active clusters in the MG stage) relative to the single peak in SRCs observed in the A^n^ subgenome in both the GLOB (53 clusters) and MG (176 clusters) seed stages. This is observed despite both regions maintaining high gene density regardless of the RE accumulation in the C^n^ subgenome relative to the A^n^ subgenome. The highly repetitive centromeric region of A06 and C06 exhibits a decline in siRNA density but is more pronounced in C06. Further, while siRNA density does increase with distal proximity to the centromere, this is especially pronounced in C06, wherein siRNA density jumps to 78 and 212 in the GLOB and MG stages, respectively (block 3, Figure 9). This increase in diversity is not seen in the corresponding syntenic blocks of A06 – the smaller syntenic blocks (block 2 and 3, Figure 9) do not accumulate SRCs as seen in C06. Further, the same syntenic blocks undergo significant changes in CG methylation in the A^n^ subgenome, but are absent in the C^n^ subgenome, while both blocks 2 and 3 experienced greater CHG methylation changes between morphogenesis and maturation in C06 than seen in A06.On the global genomic scale, we find that greater length disparity of syntenic regions does not accumulate siRNA density linearly (Dataset S10, Figure S5). In all five seed stages, syntenic blocks did not increase siRNAs to scale, even in blocks that were 2-9x larger in the C^n^ subgenome than the A^n^ subgenome. Interestingly, syntenic blocks of similar size had greater differences in SRC abundance than those with greater length disparity. This implies that the number of SRCs are conserved amongst syntenic regions and do not accumulate in longer intergenic regions. On the global genomic scale, we find that greater length disparity of syntenic regions does not accumulate siRNA density linearly (Dataset S10, Figure S5).

## Discussion

### The epigenetic bias of the C^n^ subgenome of *Brassica napus*

The genome of *B. napus* comprises relatively intact subgenomes derived from the interspecific hybridization of *B. rapa* and *B. oleracea*. Genome bias often leads to disruption of genomic integrity, wherein allopolyploid plants can undergo chromosome fractionation as bias develops (Kellogg, 2003; Cheng et al., 2018). Genome fractionation in allopolyploids occurs at a stable rate, and often involves duplicate gene loss within the submissive genome (Emery et al., 2018). The epigenome provides an additional perspective on genome fractionation and bias where the selective silencing of one genome via siRNA activity and DNA methylation may reflect incipient genome dominance. Epigenome bias is prevalent across seed development in *B. napus* where consistent bias of cytosine methylation and siRNA clusters in the C^n^ subgenome implies the selective silencing of its components, though to what extent remains uncertain. In synthesized allotetraploids interspecific hybridization can cause immediate silencing of genomic components, and lead to morphological dominance of one progenitor species over the other (Alexander-Webber et al., 2016; Comai et al., 2000; Wang et al., 2006).

Patterns of cytosine methylation and diversity of siRNA expression derived from TEs undergo substantial modifications upon allopolyploidization events. Among these, SRCs are become deregulated upon allopolyploidization and permit an increase in TE activity thereby generating instability within the genome (Kenan-Eichler et al., 2011). Cytosine methylation and siRNA diversity are biased towards the C^n^ subgenome in the *B. napus* seed indicating repression of its relative contribution to seed development and by extension is submissive to the A^n^ subgenome. These SRCs were also subject to dramatic CHH hypermethylation. This phenomenon of C^n^ subgenome bias persists within the observed increase in siRNA diversity at the MG stage where rasiRNAs are also most abundant. The greater accumulation of non-coding elements such as siRNAs and REs in the C^n^ subgenome may contribute to a greater overall increase in mCHH of the same C^n^ subgenome as the seed develops. Extensive methylation of promoters in the C^n^ subgenome and methylation on SRCs in broad syntenic blocks furthers this bias. Together, the C^n^ subgenome appears to accumulate silencing elements that persist across all stages of the seed lifecycle, and experiences selective silencing in syntenic regions corresponding to the A^n^ subgenome. As a result, syntenic regions do not mirror epigenetic structure in the *B. napus* seed. Acute subgenome bias defines the epigenetic dynamics at play during seed development in this amphidiploid species.

### The contribution of repetitive elements to subgenome bias in *B. napus* seed development

Maturation in seed development is characterized by the rapid deposition of protein and lipid energy stores for the developing seed and coincides with an enormous influx of both siRNA clusters and DNA methylation, especially in the CHH context. The influx of active siRNA clusters in the MG stage can be largely attributed to the profound increase in rasiRNA diversity. This accumulation of siRNAs derived from REs in the genome implies their importance in modulating DNA methylation, particularly in the CHH context, late in seed development. The lack of rasiRNA diversity in the DS stage is likely a product of developmental stasis achieved in seed dormancy.

Repetitive elements can be targets of epigenetic modification especially in allopolyploid plants, like *B. napus* (Parisod et al., 2010, 2009). An abundance of REs in one progenitor genome may lead to downstream epigenetic modification in a hybrid polyploid (Wendel et al., 2018). Here, we observe subgenome bias was most dramatic on REs in *B. napus* seeds and leaves. In the current study, REs are more highly methylated in the *B. oleracea* subgenome of *B. napus*. While this phenomenon has been observed in vegetative tissues (Chalhoub et al., 2014) and microspores (Li et al., 2016), our data suggests epigenetic modifications go beyond what is observed in vegetative leaves via CHH hypermethylation on REs. DNA methylation may provide an additional level of epigenetic regulation that serves to modulate variable or novel regions of the genome. Thus, trends in subgenome bias favouring higher methylation of the C^n^ subgenome may be a consequence of its higher RE load.

In *B. napus*, our data provides evidence of a global loss of methylation on REs, especially from the A^n^ subgenome, in the CG and CHG contexts relative to leaves. Thus, a higher level of methylation is still maintained on the REs in the C^n^ subgenome. Currently, we cannot detect whether this loss of methylation on REs is via a passive or active process; it is possible that loss of mCG and mCHG in seeds is a passive process that occurs over the numerous cell divisions and DNA replication cycles that establish the seed. Whether a loss of methylation relative to leaves is actively or passively achieved, it appears that C^n^ subgenome bias is a feature that persists over the transition of the plant to the next sporophytic generation.

### RdDM as a modulator of developmental transitions in seed development

Active demethylation of REs in ephemeral reproductive structures such as vegetative nuclei in pollen and the endosperm may lead to the activation of small RNA machinery to modify DNA methylation in other cells (Gehring et al. 2009, Slotkin et al. 2009, Century et al. 2012). It is known that DNA hypomethylation in *Arabidopsis* gametophytes cause alterations in seed size, and that interference in DNA methylation machinery causes loss of seed viability (Zhang et al., 2010; Xiao et al., 2006b, 2006a). Interestingly, the two stages exhibiting the highest siRNA diversity in our dataset were the OV and MG stage. The bimodal accumulation of siRNAs at gametogenesis and again at maturation fall in line with the developmental importance of these stages that lead to the initiation of the seed and the finalization of its development prior to dormancy. Given the predominant role siRNAs play in managing DNA methylation via the canonical RdDM pathway, the siRNA diversity early and late in seed development may indicate versatile processes vital to seed development depend on the action of siRNAs in *B. napus.* Patterns of siRNA accumulation and cytosine methylation shift around genes involved in cell division and growth in *B. napus* seed development. During early seed development, siRNAs accumulate in promoters of genes associated with zygote and endosperm development, whereas in late seed development, promoters of genes involved in cell division and auxin/giberellin/cytokinin are targeted for CHH hypermethylation. These targets reflect the developmental phenotypes observed in Arabidopsis lacking DNA methylation machinery, wherein improper cellular divisions are frequently observed (Kim et al., 2008; Xiao et al., 2006b, 2006a).

RdDM in maternal tissues is essential for early *B. rapa* seed development (Grover et al. 2018). Given that the maternal seed coat makes up a substantial portion of the *B. napus* seed at this stage, it is possible that some patterns of early RdDM activation is maintained from its progenitor *B. rapa*. The role of RdDM in seed maturation is still unclear, though it appears to be characteristic of seed maturation, but viably non-essential in the model system Arabidopsis (Lin et al., 2017; Bouyer et al., 2017). We found that siRNA loci were heavily methylated in mature *B. napus* seeds, where the embryo comprises the vast majority of the seed body. A spatiotemporal analysis of DNA methylation and small RNA accumulation in the embryo, endosperm, and seed coat would provide insight into the RdDM in the maternal and zygotic tissues throughout seed development, and whether the embryo, endosperm, and seed coat are all subject to the same regimes of subgenome bias observed in whole seeds.

### Subgenome bias of rasiRNAs is progressively lost throughout seed development

Early in seed development rasiRNAs constitute much less of the total siRNA profile when compared to maturation stages. Despite this, the Cn subgenome bias of rasiRNA species are most accentuated early in seed development and is progressively lost as the seed matures. This suggests the dominant action of the A^n^ subgenome is most prevalent early in seed development, with the C^n^ subgenome being asymmetrically silenced by the action of rasiRNAs during gametogenesis and morphogenesis.

Maturation in seed development is characterized by the rapid deposition of protein and lipid energy stores for the developing seed and coincides with an enormous influx of both siRNA clusters and DNA methylation, especially in the CHH context. The influx of active siRNA clusters in the MG stage can be largely attributed to the increase in rasiRNA diversity. This accumulation of siRNAs derived from REs in the genome implies their importance in modulating DNA methylation, particularly in the CHH context, late in seed development. The erasure of rasiRNA diversity in the DS stage is likely a product of developmental stasis achieved in dormancy. This peak in diversity is complemented by the lowest bias in the C^n^ subgenome and extensive global CHH methylation. This is echoed by extensive CHH methylation observed during seed maturation in the diploid model plant *Arabidopsis* (Kawakatsu et al., 2017), and suggests that the global CHH methylation characteristic of maturation overrides subgenome bias in allopolyploids.

### phasiRNA diversity is abolished in seed maturation of *B. napus*

The presence of phasiRNAs in reproductive tissues is most extensively documented in the Poaceae but is now emerging to be important in dicot lineages (Xia et al., 2019). It is known that phasiRNAs are implicated in seed coat development and flowering time in the Malvaceae and Rutaceae, respectively, but reproductive 24-nt phasiRNAs are thought to be broadly absent in the Brassicaceae (Liu et al., 2017; Zhao et al., 2020) Despite this, the OV, GLOB, and HEART stages accumulate the vast majority of phasiRNAs in *B. napus* seed development and may be reflective of the reproductive roles phasiRNAs perform in the aforementioned taxa. Here, we report that 24-nt phasiRNAs accumulate during early reproductive stages of seed development and become depauperate as the seed matures. The restricted window of phasiRNA activity in *B. napus* may account for the lack of documentation of the 24-nt phasiRNA pathway in the Brassicaceae. This was predicted by Xia et al., (2019), and suggests the pathway exists in the family, though it may have only been retained in lineages outside the Camelineae.

### Synteny as an important predictor of genome bias in allopolyploids

It is known that even in extensively polyploid clades that syntenic blocks are preserved throughout generations of ploidy changes and genomic stress (Hardigan et al., 2020). However, while stability of syntenic regions may be preserved over extensive evolutionary time in polyploids, it is unknown whether the epigenome of syntenic regions in allopolyploid progenitor genomes experience the same retention. The epigenome is vitally important to the management of genome stability and provides a provisional solution to times of genomic shock (Ha et al., 2009; Springer et al., 2015). We find evidence for the lens of synteny to be valuable in demonstrating bias at the epigenome level, wherein syntenic regions across the genome experience asymmetric DNA methylation and accumulation of siRNAs not seen globally in the genome. As it stands, the variables determining which subgenome persists as dominant are still enigmatic, but the epigenetic biases documented here in the nascent amphidiploid *B. napus* suggest epigenomic bias begins and establishes early in the species’ history. As a result, epigenome architecture of syntenic regions may be an important factor in determining the evolutionary trajectory of the species and is especially relevant to the ubiquitously polyploid crop plants.

## Materials and Methods

The DH12075 genome sequence and annotation were provided by Drs. Isobel Parkin and Steve Robinson via the Canadian Canola Genome Sequencing industry consortium project.

### Plant growth conditions and sample collection

Methylome analyses were performed in B. napus cv. DH12075. Plants were grown in growth chambers under long day conditions at 22°C. Flowers were hand-pollinated and collected at 0 DPF (OV), 7 DPF (GLOB), 10 DPF (HEART), 28 DPF (MG), and 35 DPF (DS). Seeds were manually harvested from siliques and ground in liquid nitrogen for nucleic acid extractions. DNA was isolated from seeds using the Qiagen Plant DNeasy Mini Kit. RNA was isolated from seeds using the Qiagen miRNeasy mini kit. Three bioreplicates were used for WGBS and sRNAseq.

### Synteny map of the DH12075 genome

Syntenic regions were determined using MCScan under default parameters (Tang et al., 2008). The DH12075 A^n^ subgenome was compared to the C^n^ subgenome to output the colinear regions linking the two.

### Methylome library preparation and sequencing

Library preparation for bisulfite sequencing was performed at Genome Quebec. DNA was sheared via sonication on the Covaris platform prior adaptor ligation and bisulfite conversion using the EZ-96 DNA Methylation-Lightning Kit (Zymo®). Libraries were then amplified using the DNA UltraII Kit (NEB®). Libraries were sequenced on the HiSeq X platform (2 lanes, 150bp PE). Bisulfite-sequencing was performed on three replicates for both stages. A total of 1.24 billion 150bp paired-end reads were generated from the globular and mature green libraries. Chloroplast genome cytosine methylation was used as a proxy to estimate bisulfite non-conversion rate -conversion efficiency is estimated to be ~98%. Bisulfite-sequencing libraries (> 97% bisulfite conversion rate) (Dataset S5) for both samples. Approximately 67% of paired reads aligned to the *B. napus* cv. DH12075 genome (Dataset S5). Differential methylation was determined over 400bp windows using MethylKit (Akalin et al. 2012). Promoter regions were calculated as 1000bp regions upstream of the TSS of a gene. To identify genes with upstream regulatory regions containing DMRs, overlap of the 400bp DMR regions and the 1000bp upstream promoter regions was calculated using BEDtools. Homologous genes in the Darmor Bzh annotation were identified, and enrichment of transcript accumulation patterns was performed using SeqEnrich (Becker et al. 2017).

### Bisulfite sequencing analysis

Quality trimming and adaptor removal were performed using Trimmomatic (http://www.bioinformatics.babraham.ac.uk/projects/fastqc/). Successfully paired reads were aligned to the DH12075 genome using BSMap2, and cytosine methylation levels were called using the methratio script (Xi and Li 2009). Individual cytosines had to be covered by at least 4 reads to be considered for analysis. Average DNA methylation over 1kB regions, and average methylation ratios of genes, REs, and promoters were calculated using BEDtools (Quinlan and Hall 2010). We also examined the effect of bin size on determining subgenome bias in average methylation and found that the C subgenome was detected as more highly methylated regardless of bin size (Dataset S5). Differential methylation was determined over 400bp windows using MethylKit (Akalin et al. 2012). Promoter regions were calculated as 1000bp regions upstream of the TSS of a gene.

### Small RNA library preparation and sequencing

Total RNA collected from seeds was used for library prep with the NEBNext^®^ Small RNA Library Prep Set for Illumina^®^ (Multiplex compatible). Libraries were pooled and sequenced on the Illumina HiSeq2500 (2 lanes, 50bp SE). Quality trimming and adaptor removal were completed with Trimmomatic (Dataset S5). Surviving reads were aligned to the DH12075 genome using Bowtie on default settings, with an average of 87% alignment (Dataset S5). Aligned reads were then filtered for rRNA, tRNA, snRNA, and snoRNA species by aligning them to known records in NCBI under all Viridiplantae.

### SRC identification and analysis

siRNA prediction was done using Short Stack on default settings (Axtell, 2013). SRCs overlapping 90% with a predicted SRC in another bioreplicate were passed through analysis to coincide with miRNA criteria covered in Axtell and Meyers (2018). SRCs overlapping with REs were classified as rasiRNAs, and SRCs exceeding a phase score of 25 were classified as phasiRNAs. siRNA density was calculated in 100 kB windows. Homologous genes in the Darmor Bzh annotation were identified and GO enrichment on genes downstream of SRCs localized to the corresponding promoter was performed using SeqEnrich (Becker et al. 2017).

### GEO Accession

Note: at the time of submitting this manuscript, our data was still uploading to NCBI. The GEO accession number should be available to the reviewers at this time.

## Supplemental Data Files

Dataset S1. Syntenic block coordinates and short list of the longest contiguous syntenic block pair per chromosome. Average methylation within each of the largest syntenic blocks included both as a table and as a scatterplot visualization.

Dataset S2. Methylation levels over 1kB bins for leaves, globular stage seeds (GLOB), and mature green stage seeds (MG). Methylation levels of all genes, promoters, and transposable elements are also included. Mean methylation levels were calculated based on individual cytosine methylation levels in each 1kB bin using BEDtools. Each context was considered separately. Methylation levels calculated across different bin sizes are also shown.

Dataset S3. Cytosine methylation over the largest syntenic blocks. Average methylation is shown for CG, CHG, and CHH contexts for leaves, globular (GLOB), and mature green (MG) seeds over 1kb windows, genes, promoters, and repetitive elements (REs) that overlapped the largest syntenic regions between the An and Cn subgenomes. The results of the Mann-Whitney Wilcox test (Bonferroni correction) are included.

Dataset S4. Methylation levels over 1kB bins for leaves, globular stage seeds (GLOB), and mature green stage seeds (MG). Methylation levels of all genes, promoters, and transposable elements are also included. Mean methylation levels were calculated based on individual cytosine methylation levels in each 1kB bin using BEDtools. Each context was considered separately. Methylation levels calculated across different bin sizes are also shown.

Dataset S5. Alignment rates and QC of WGBS and sRNAseq data

Dataset S6. Complete identifier (Chromosome_start_stop) and sequence of all siRNA clusters (sequences are of complete cluster, and do not necessarily indicate the mature siRNA sequence) in each developmental stage: OV, GLOB, HEART, MG, DS.

Dataset S7. siRNA clusters identified as either rasiRNAs or phasiRNAs in each developmental stage: OV, GLOB, HEART, MG, DS. siRNAs intersecting with repetitive elements deemed rasiRNAs, and siRNAs exceeding a phase score of 25 per Short Stack’s analysis were deemed phasiRNAs.

Dataset S8. Total number of SRCs intersecting with genes and promoters. Complete gene list of SRCs intersecting with promoters, and the GO enrichment of the same gene lists.

Dataset S9. siRNA cluster density in 1000kB windows in each developmental stage: OV, GLOB, HEART, MG, DS.

Dataset S10. Length and siRNA cluster quotient of the C and A subgenomes, respectively. Scatterplots are present to visualize the relationshop between the degree of genomic bias in length and in siRNA abundance within syntenic pairs.

## Author Contributions

D.K. and D.J.Z. designed and performed research, analyzed data, and co-wrote the article. N.P.V. developed data analysis tools, drafted figures, and analyzed data. I.A.P. and S.J.R. contributed the DH12075 genome build/annotation and provided feedback on the manuscript. M.F.B. designed research and co-wrote the article.

## Acknowledgements

We would like to thank Dr. Olivia Wilkins for their feedback and helpful discussion of the manuscript. We would also like to thank Grigory Shamov and Ali Karrache (WestGrid/Compute Canada) for their technical support. Sequencing was performed at Genome Québec. This work was supported through generous funding from the Natural Science and Engineering Research Council of Canada to Mark Belmonte.

---

**Figure S1.**
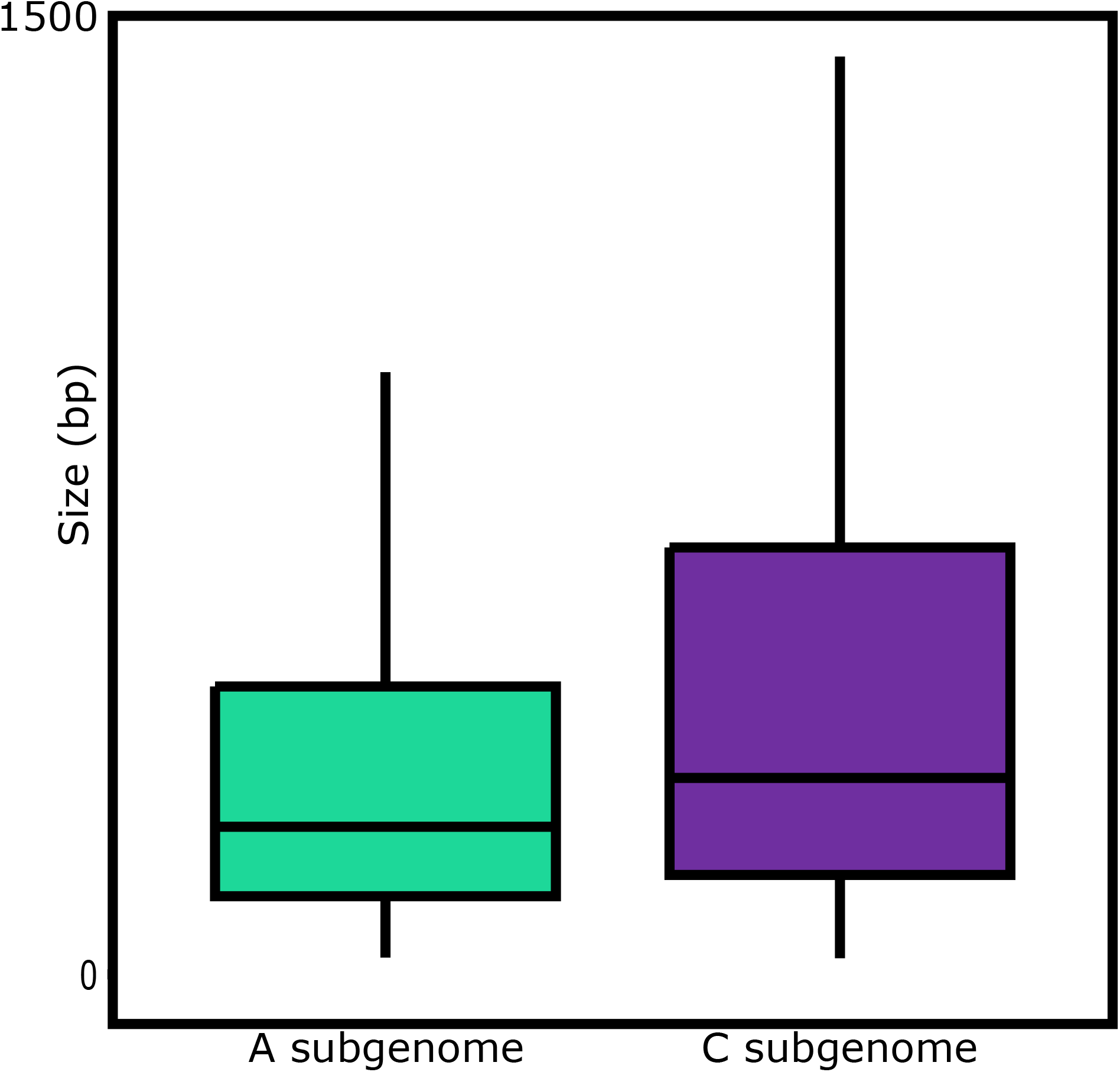
Quantile boxplots showing the distribution of size of repetitive elements (REs) in the A^n^ (teal) and C^n^ (purple) subgenomes of *B. napus* cv. DH12075.

**Figure S2.**
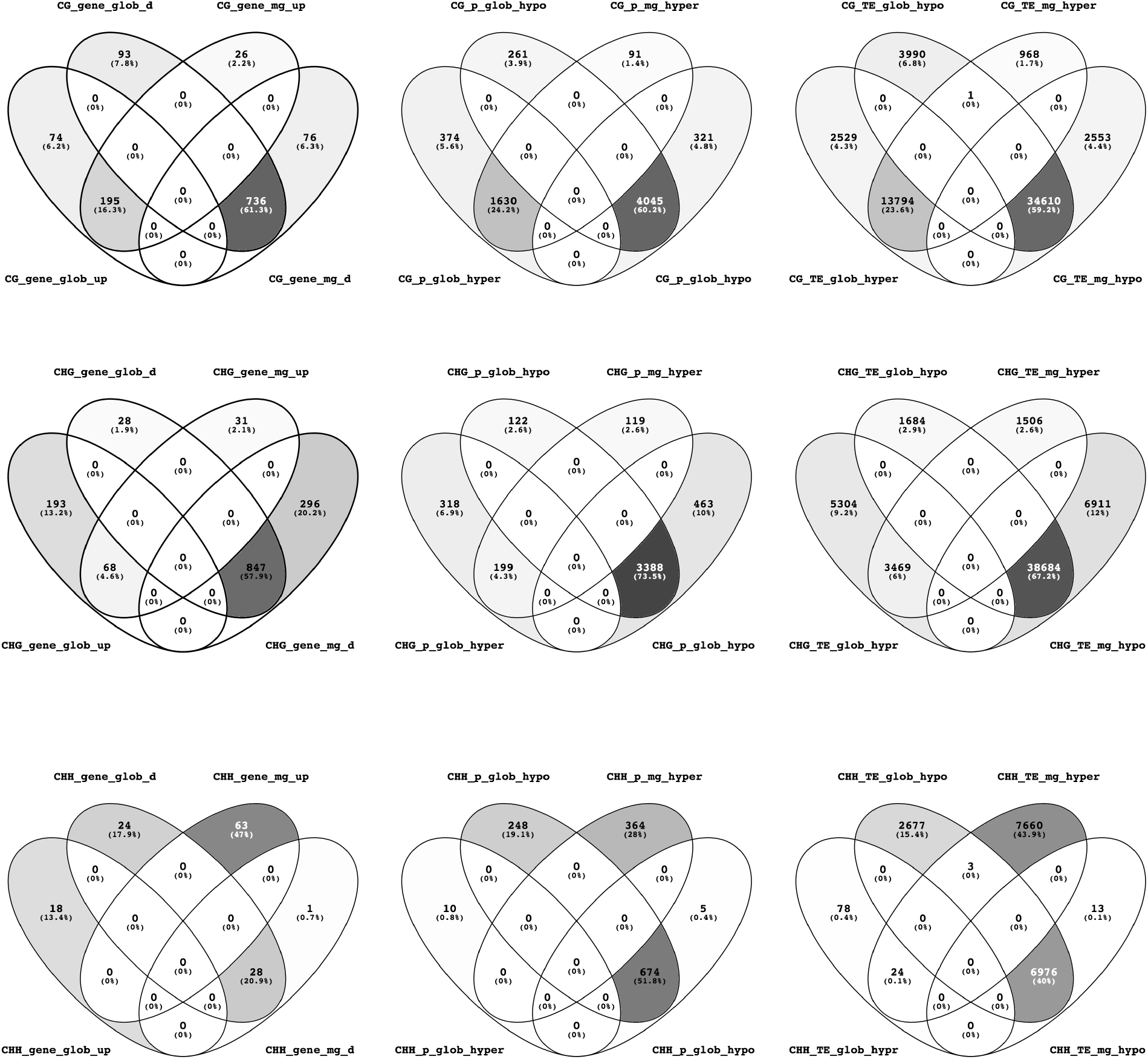
Differential methylation between leaves and seeds. Venn diagrams showing hyper and hypo methylated genes, promoters and repetitive elements in GLOB and MG seeds relative to leaves. Differential methylation was determined using MethylKit over 400bp windows of the genome. Intersection with transposable elements was determined using BEDtools. Gene lists for Venn intersects can be found in Data DK3.

**Figure S3.**
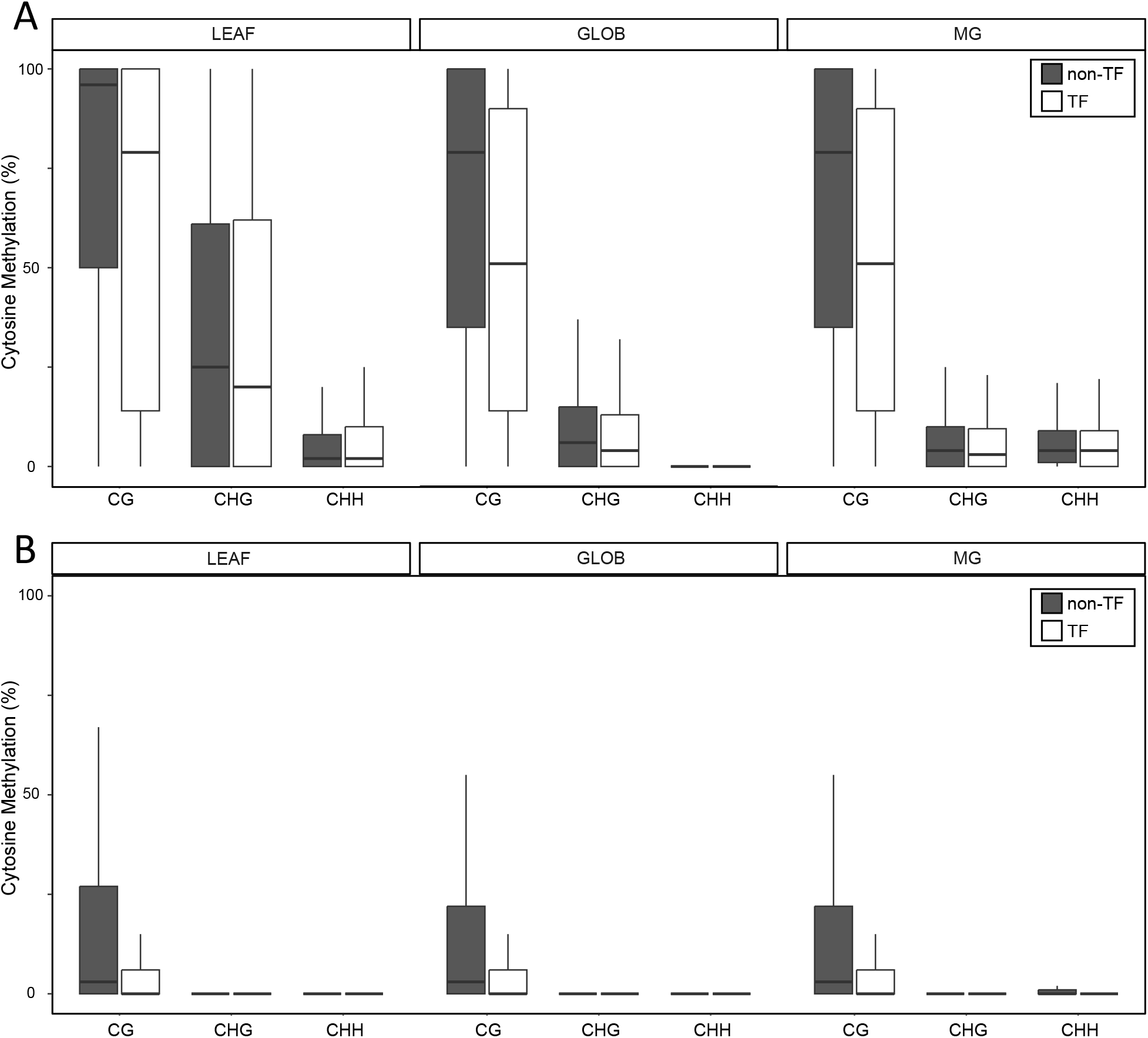
Methylation levels of transcription factors. Quantile boxplots illustrating the methylation levels of gene bodies (A) and promoters (B; 1kb upstream of TSS) of non-transcription factor genes (grey) and transcription factors (white) in the CG, CHG, and CHH contexts in leaves, and globular (GLOB) and mature green (MG) seeds).

**Figure S4.**
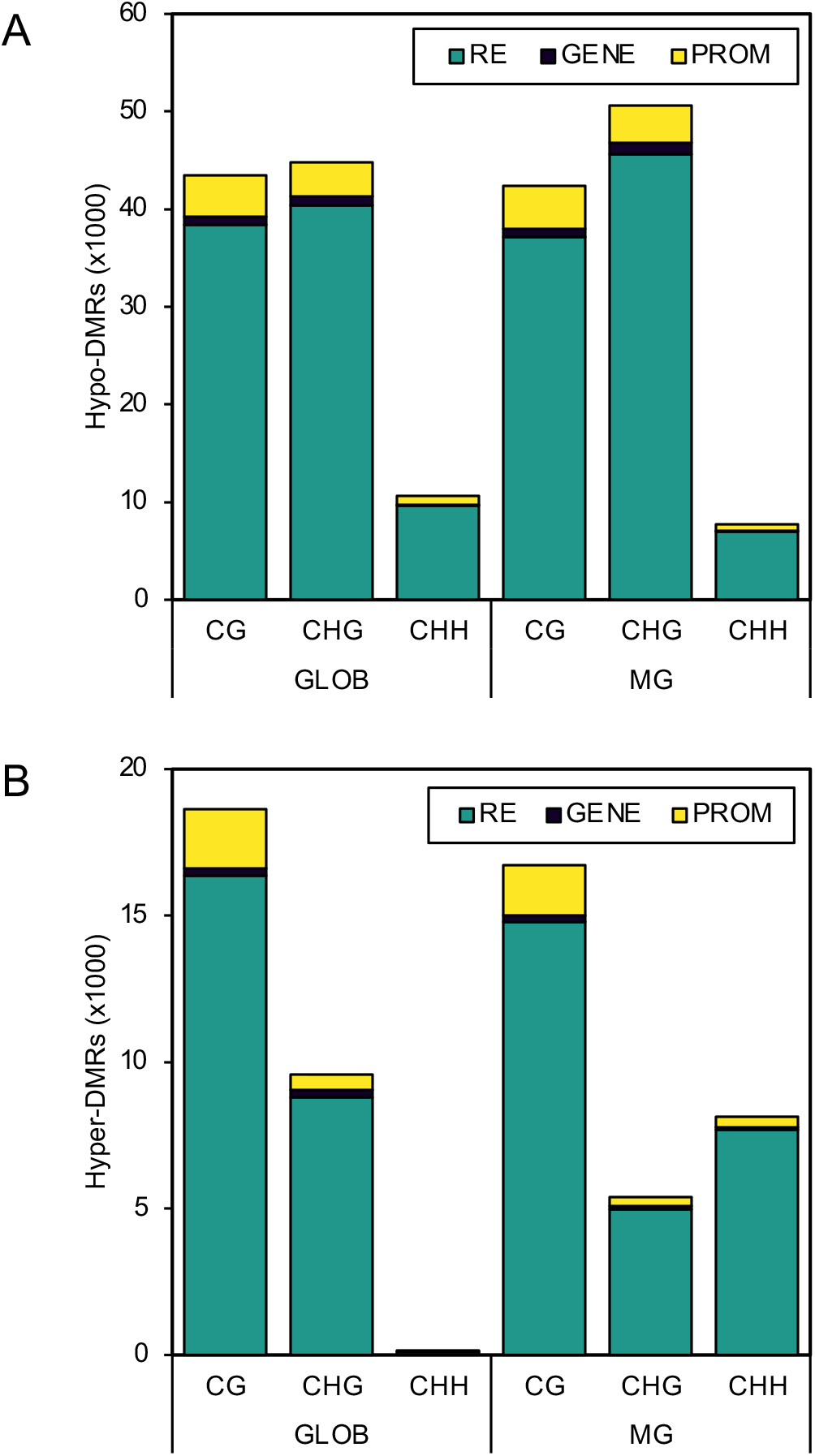
Shifts in DNA methylation between in development. (A-B) Differential cytosine methylation in seeds relative to leaves. The GLOB and MG seed methylomes were compared to the leaf methylome (Parkin et al., unpublished). Differential methylation was calculated for all three cytosine contexts (CG, CHG, CHH) over genes, promoters and repetitive elements (REs). Number of differentially methylated REs (teal), gene bodies (purple) and promoters (yellow) that are (A) hypomethylated in the GLOB or MG seed relative to leaves or (B) hypermethylated in the GLOB or MG seed relative to leaves.

**Figure S5.**
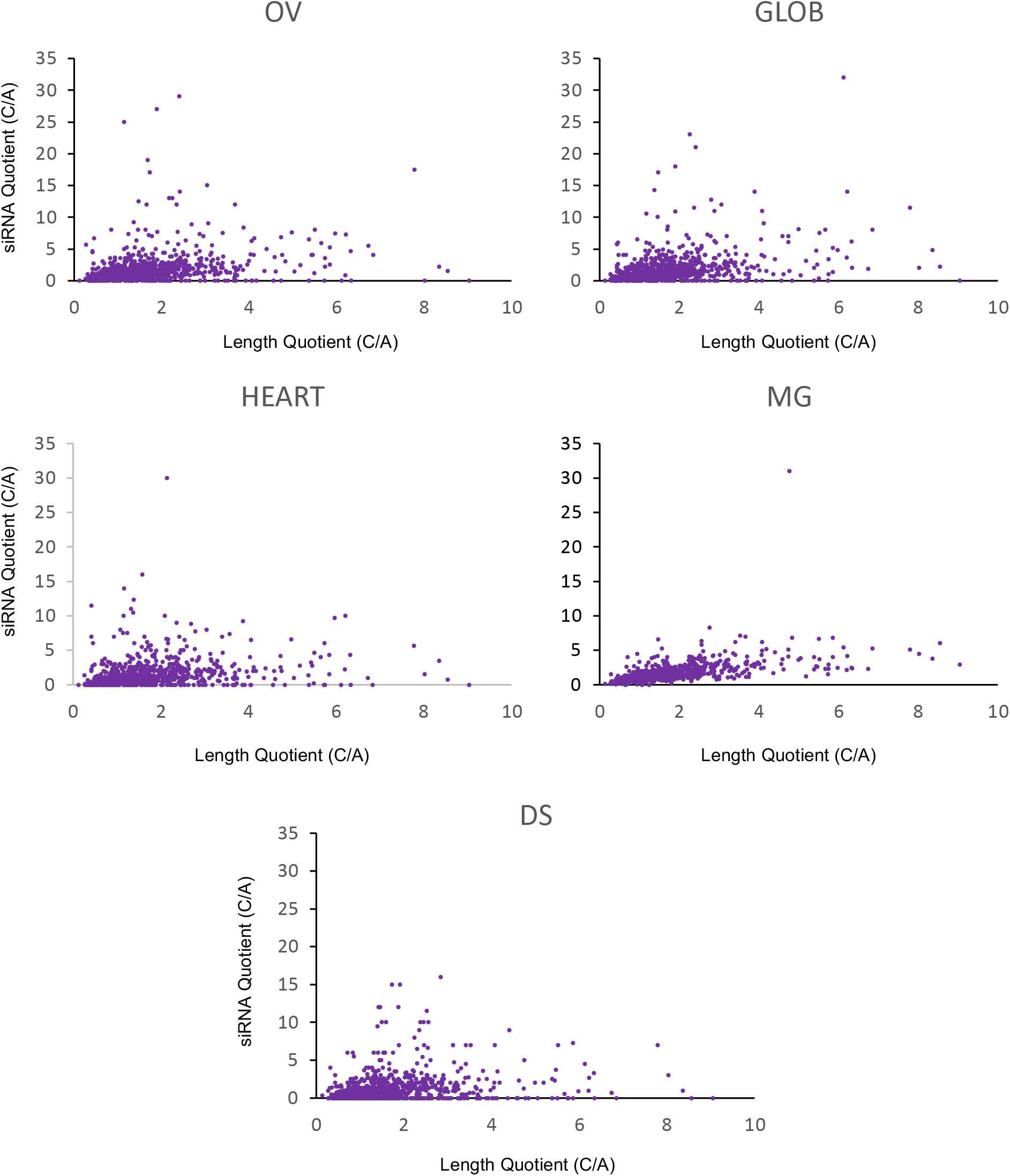
Relationship between length disparity of syntenic regions and siRNA abundance between the A^n^ and C^n^ subgenomes. siRNA quotient represented by total siRNA clusters intersecting with the syntenic block of the C^n^ subgenome divided by the number of siRNA clusters in the corresponding syntenic block in the A^n^ subgenome. Length quotient represented as the length of the C^n^ subgenome’s syntenic block length (in bp) divided by the length of the A^n^ subgenome’s syntenic block length (in bp). siRNAs active in seed development do not accumulate proportionately with high disparity in syntenic block length.

